# MemConverter: An Iterative Pipeline for Reprogramming Protein Localization in Membrane or Aqueous Solution

**DOI:** 10.1101/2025.10.23.684164

**Authors:** Jun Li, Haozhe Guo, Chen Song

## Abstract

We propose a pipeline, MemConverter, for the conversion of soluble proteins to membrane proteins and vice versa, based on the prediction of membrane contact probability (MCP). By fine-tuning ProteinMPNN with a membrane protein dataset to create MemProtMPNN, and integrating it with iterative structure refinement using AlphaFold2, our approach enables precise reprogramming of protein surface properties. The pipeline utilizes MCP-guided sequence fusion to direct the evolution of protein sequences towards the desired membrane or soluble environments. During this process, selective residues are fixed to concentrate design efforts on regions requiring alteration. The pipeline demonstrates superior performance compared to existing methods, and molecular dynamics simulations validate that the designed proteins exhibit stable membrane integration or enhanced solubility, depending on the target localization. This pipeline introduces a novel computational tool for engineering proteins to be localized in membranes or aqueous solutions.

## 1 Introduction

Membrane proteins play a pivotal role in cellular processes, including signal transduction^1^ and molecular transport.^2^ Consequently, a comprehensive understanding of their structures, properties, and functions is essential for investigating related biological processes and for designing novel membrane proteins. Building on existing knowledge of proteins, the field of protein design has advanced significantly in recent years, catalyzed by a new generation of AI-based computational tools. Methods for protein structure design, such as RFDiffusion, ^3^ FrameDiff, ^4^ Genie ^5^ and TopoDiff, ^6^ along with methods for protein sequence design, like ProteinMPNN, ^7^ ESM-IF, ^8^ and CarbonDesign, ^9^ have dramatically accelerated the protein design process. However, the *de novo* design of membrane proteins remains a significant challenge, ^10^ with only a limited number of successful cases. ^11–15^

Due to the significant challenges in expressing, purifying, and functionally validating membrane proteins, their *de novo* design often employs a *soluble-first* strategy. ^13,15^ In this paradigm, a soluble protein analogue with the desired function is first created and validated in an aqueous environment, and is subsequently transformed into an integral membrane protein by modifying its surface residues to be more hydrophobic. This transformation process requires precise delineation of the protein’s transmembrane regions and strategic substitution of amino acids to maintain function. Although conventional methods based on Rosetta have achieved notable successes, ^16^ their heavy reliance on the quality of the initial structure, the accuracy of regional demarcation, and the designer’s expertise has limited the advancement of functional membrane protein design. Recently, a deep learning-based method was reported for effectively converting a designed water-soluble fluorescent activating protein (wFAP) into a functional transmembrane protein (tmFAP) using gradient-guided hallucination. ^15^ Nevertheless, this approach still depends on predefined amino acid composition patterns across different structural regions.

On the other hand, for the purpose of studying existing membrane proteins rather than *de novo* designing them, a reverse strategy is often employed. A key objective in this context is to simplify experimental characterization by engineering membrane proteins into water-soluble analogues. This is achieved by substituting hydrophobic residues on the transmembrane surface with hydrophilic ones, which facilitates extraction and purification while preserving the core structural and functional properties. ^15^ However, a systematic and automated approach for such conversion is not yet available.

Recently, we proposed a characteristic quantity (or metric), *membrane contact probability*, ^17,18^ to quantify the probability of residues in a given protein sequence being in direct contact with the hydrophobic cores of membranes. The latest version of the MCP predictor, ProtRAP-LM, which leverages protein language models, has significantly improved the accuracy and speed of MCP predictions. ^19^ Since this deep learning-based model has learned the intrinsic features of membrane protein sequences, we believe that the predicted MCP can be utilized to refine existing protein sequence generators without requiring input from human experts, thereby enhancing eficiency in the design of membrane proteins.

Additionally, it has been recognized that subtle yet essential structural distinctions may exist between integral membrane proteins and their corresponding soluble analogues. According to the study by Goverde et al. on designing soluble analogues of membrane proteins, certain structural differences between these protein types have been identified.^20^ Furthermore, as demonstrated in the de novo design of ion channels from water-soluble *α*-helical barrels, the conversion from soluble to membrane-embedded forms can induce notable topological rearrangements. ^13^ However, for a given monomeric protein, these discrepancies typically manifest as subtle differences, which arise from adaptations that reduce transmembrane hydrophobicity for aqueous solubility while preserving core topologies. ^20^ We therefore contend that controlled structural perturbations during the bidirectional conversion process are indispensable for the desired localization design.

Therefore, in this study, we present an MCP-guided pipeline, MemConverter, to convert soluble analogues into integral membrane proteins and vice versa. To achieve this, we have fine-tuned ProteinMPNN, developing a specialized membrane protein sequence generator termed *MemProtMPNN*. The MemConverter pipeline utilizes MemProtMPNN and SolubleMPNN ^20^ to iteratively optimize the protein sequence, followed by the application of AlphaFold2 for structural refinement. This process facilitates the gradual conversion of soluble proteins to membrane proteins and the reverse, guided by the predicted MCP. The localization of the designed proteins in either membrane or aqueous solution is validated *in silico* through molecular dynamics (MD) simulations.

## 2 Results

### 2.1 MemProtMPNN for sequence generation of membrane proteins

To convert soluble proteins into membrane proteins, it is essential to utilize a sequence generation model specifically designed for membrane proteins. ProteinMPNN is the most widely utilized foundation model for sequence generation. Based on this, while SolubleMPNN has been proposed for the design of soluble proteins, ^20^ to the best of our knowledge, there are few models that have been specifically tailored for the design of membrane protein sequences. To address this gap, we developed MemProtMPNN to facilitate robust sequence design for membrane proteins.

Rather than training a model from scratch, we leveraged the inherent capabilities of ProteinMPNN, which represents the state of the art in mapping protein structure to sequence. We fine-tuned ProteinMPNN using the isolated transmembrane domain dataset extracted from tmAFDB. ^21^ More specific details regarding the dataset construction and training methods are included in the Methods section.

To evaluate the performance of MemProtMPNN in comparison to ProteinMPNN for membrane protein sequence design, we adopted a series of metrics (details in the Methods section). We first analyzed the distribution of the pLDDT score of the designed sequences as a function of transmembrane domain length. As illustrated in Fig. 1a, for proteins ranging in length from 100 to 450 residues, MemProtMPNN demonstrates a significant performance improvement compared to the standard ProteinMPNN. Furthermore, Fig. 1b presents the distribution of pLDDT scores, with points skewed toward the upper-left quadrant indicating a modest but consistent overall enhancement for MemProtMPNN over ProteinMPNN. In addition, evaluations of sequence recovery rates across varying residue exposure levels revealed that MemProtMPNN maintained a consistent 5% advantage, spanning from buried to exposed residues (Fig. 1c). Successful designs were defined as those with scRMSD < 2 Å; among 3,722 tested cases, MemProtMPNN achieved 1,076 successes (28.91%), compared to 867 successes (23.29%) for ProteinMPNN. For a clearer comparison of peak model performance, the top 1,000 proteins from each model were selected and contrasted in a violin plot (Fig. 1d), showing that MemProtMPNN’s performance is indeed overall better than ProteinMPNN. We also assessed deviations in the frequencies of amino acid residues in the designed versus native membrane proteins (Fig. 1e), where MemProtMPNN exhibited superior fidelity across most residue types, particularly for hydrophilic residues such as D, C, and S. Notably, a slight reduction in sequence diversity was observed with MemProtMPNN (Fig. 1f), which likely stems from the fact that membrane proteins have a more limited structure and sequence diversity than soluble proteins. ^22^

**Figure 1:**
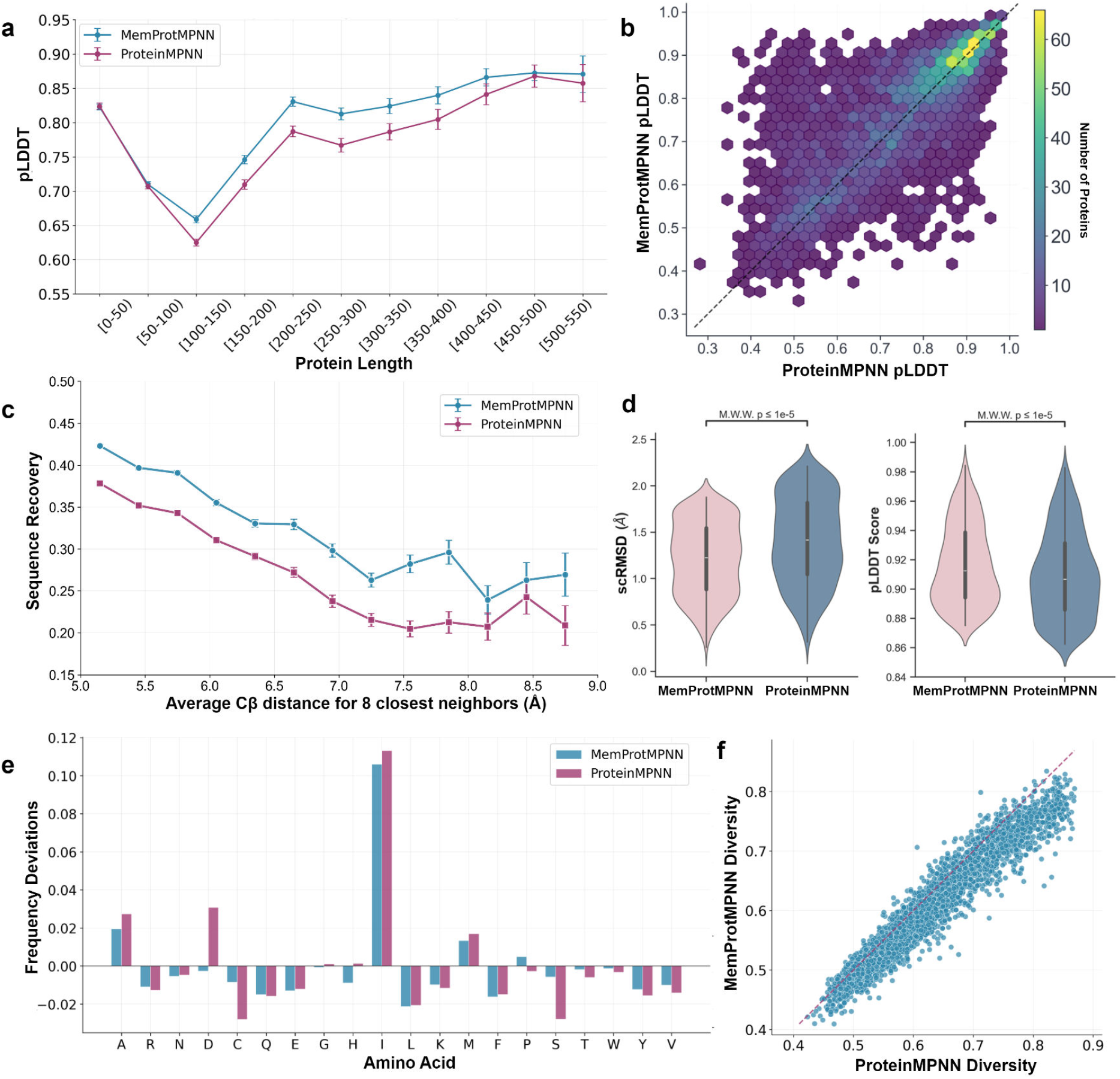
Performance comparison between MemProtMPNN and ProteinMPNN. **a** Variation of average pLDDT with protein length for designed sequences. **b** Hexbin plot showing the joint distribution of pLDDT scores for designed sequences, with ProteinMPNN predictions on the x-axis and MemProtMPNN predictions on the y-axis. **c** Comparison of sequence recovery rate across varying residue exposure levels, measured by average Cβ distance for the eight closest neighbors. **d** Comparative pLDDT and scRMSD for top 1,000 designed proteins. **e** Frequency deviations in amino acid usage between designed and native sequences. **f** Scatter plot of sequence diversity for designed sequences, comparing ProteinMPNN (x-axis) with MemProtMPNN (y-axis).

The MemProtMPNN is employed as the sequence generation module in our pipeline for the conversion of membrane proteins, as detailed in the following sections.

### 2.2 The MemConverter Pipeline

Coupling sequence generation with iterative structure refinement represents a promising new direction in protein design. The potential of this strategy was recently demonstrated by the af2cycler pipeline, ^23^ which was developed to enhance the designability of novel protein structures generated by models such as Chroma. ^24^ In this study, we adapt this iterative approach to address a distinct challenge: converting proteins between soluble and membrane environments.

Our computational pipeline is illustrated in Fig. 2. The framework operates through an iterative process. In each cycle, sequence design is coupled with structure prediction to progressively transform the protein sequence and structure. Central to this method is the MCP-guided sequence fusion step, which facilitates the gradual evolution of the sequence into membrane proteins by leveraging ProtRAP-LM’s intrinsic understanding of membrane protein sequences. ^19^ The entire workflow is detailed below.

**Figure 2:**
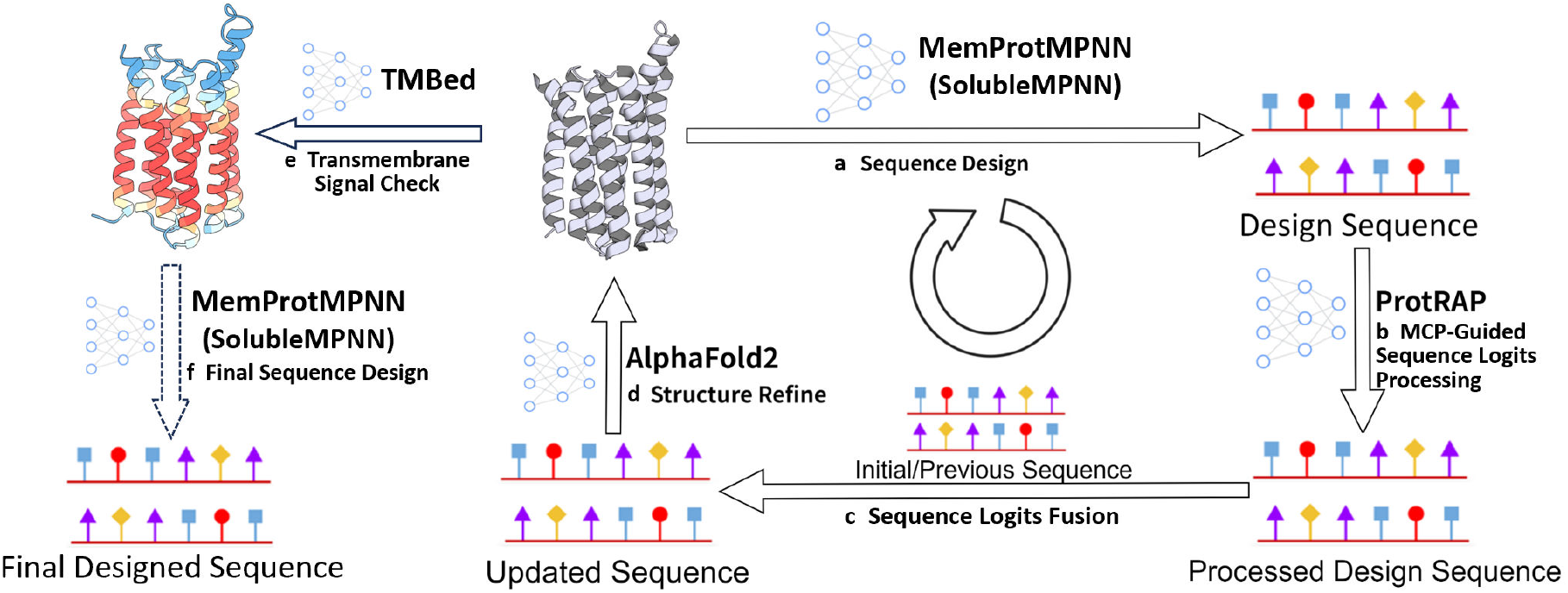
The overall framework of the MemConverter pipeline. Each iterative cycle sequentially includes steps **(a)** through **(d)**, with the loop termination criterion assessed in step **(e)**. Upon convergence and exit from the loop, the final designed sequence is generated in step **(f)**. The sequence generation module is substituted with SolubleMPNN to perform the reverse (membrane-to-soluble) conversion.

#### Sequence Design with MemProtMPNN

The pipeline begins with the structure of a known soluble protein. In each iteration, MemProtMPNN is utilized to design sequences with a high propensity for a membrane environment (Fig. 2a).

#### MCP-Guided Logits Fusion

The designed sequence from the previous step may not fully conform to the characteristics of a membrane protein; therefore, a sequence fusion step is required under the guidance of MCP. This step involves mutating residues to progressively align the sequence with the MCP profile. This process is divided into two phases, as illustrated in Fig. 2b and Fig. 2c. Through MCP-guided processing of sequence logits, low-confidence residues (determined by the discrepancy between predicted and target MCP) are partially perturbed by adding noise to explore alternatives, while high-confidence residues are retained (Fig. 2b). This preserves logits for residues with higher MCP values and perturbs those with lower values, ensuring precise control over sequence evolution toward the desired membrane-embedding characteristics. Subsequently, a weighted fusion of the post-processed logits and those from the previous (or initial) sequence is performed to promote gradual evolution (Fig. 2c). A more detailed explanation is provided in the Methods section.

#### Structure Prediction and Evaluation

After the sequence fusion, we use AlphaFold2 to predict the corresponding structure (Fig. 2d). As the input sequence is progressively optimized for a membrane environment, we anticipate that the predicted structure will be gradually refined to align more closely with the characteristics of membrane proteins.

#### Cycle and Final Candidate Selection

The design cycle is conducted iteratively. After each cycle, the redesigned sequence is evaluated using TMBed ^25^ to assess the formation of transmembrane topology. Meanwhile, the structures predicted by AlphaFold2 are evaluated using the pLDDT metric to assess sequence quality (Fig. 2e). The design cycle continues until TMBed successfully identifies a stable transmembrane signal, at which point the iterative process terminates, and the designed sequence is selected as the final candidate.

The iterative process yields a novel protein backbone optimized for a membrane environment. This final structure serves as a fixed scaffold for *de novo* sequence design using MemProtMPNN (Fig. 2f). The designed candidate sequences are evaluated based on multiple metrics, including pLDDT, scRMSD (self-consistency RMSD), and transfer free energy from solution to membrane. The best sequences can be selected for in silico or experimental validation.

#### Bidirection conversion using different sequence generation models

It should also be noted that, the sequence design model is a modular component of our framework. The custom-trained MemProtMPNN described above can be readily substituted with other sequence generation models for membrane proteins. Moreover, this modular architecture enables the reversal of the design objective; by replacing MemProtMPNN with SolubleMPNN, the pipeline can be repurposed to design soluble variants of native membrane proteins.

### 2.3 Conversion of soluble proteins to membrane proteins

#### 2.3.1 Four case studies with our pipeline

We apply our pipeline to transform soluble proteins into membrane variants across four case studies: GPCR’s soluble analogues (PDB: 8oyy), ^20^ Rhomboid Protease’s soluble analogues (PDB: 8oyw), ^20^ Claudin’s soluble analogues (PDB: 8oyv) ^20^ and a *de novo* designed water-soluble fluorescence binding protein (PDB: 8w6f). ^15^

A series of sequences was generated for each case using our pipeline, and high-quality sequences were selected based on pLDDT scores and transfer free energy. As exemplified by the 8oyy case in Fig. 3b and Fig. 3c, the soluble protein undergoes significant surface remodeling, characterized by a transition from hydrophilic to hydrophobic regions and a reduction in polar and charged residues. This remodeling results in more neutral electrostatic potentials, which facilitate membrane integration.

**Figure 3:**
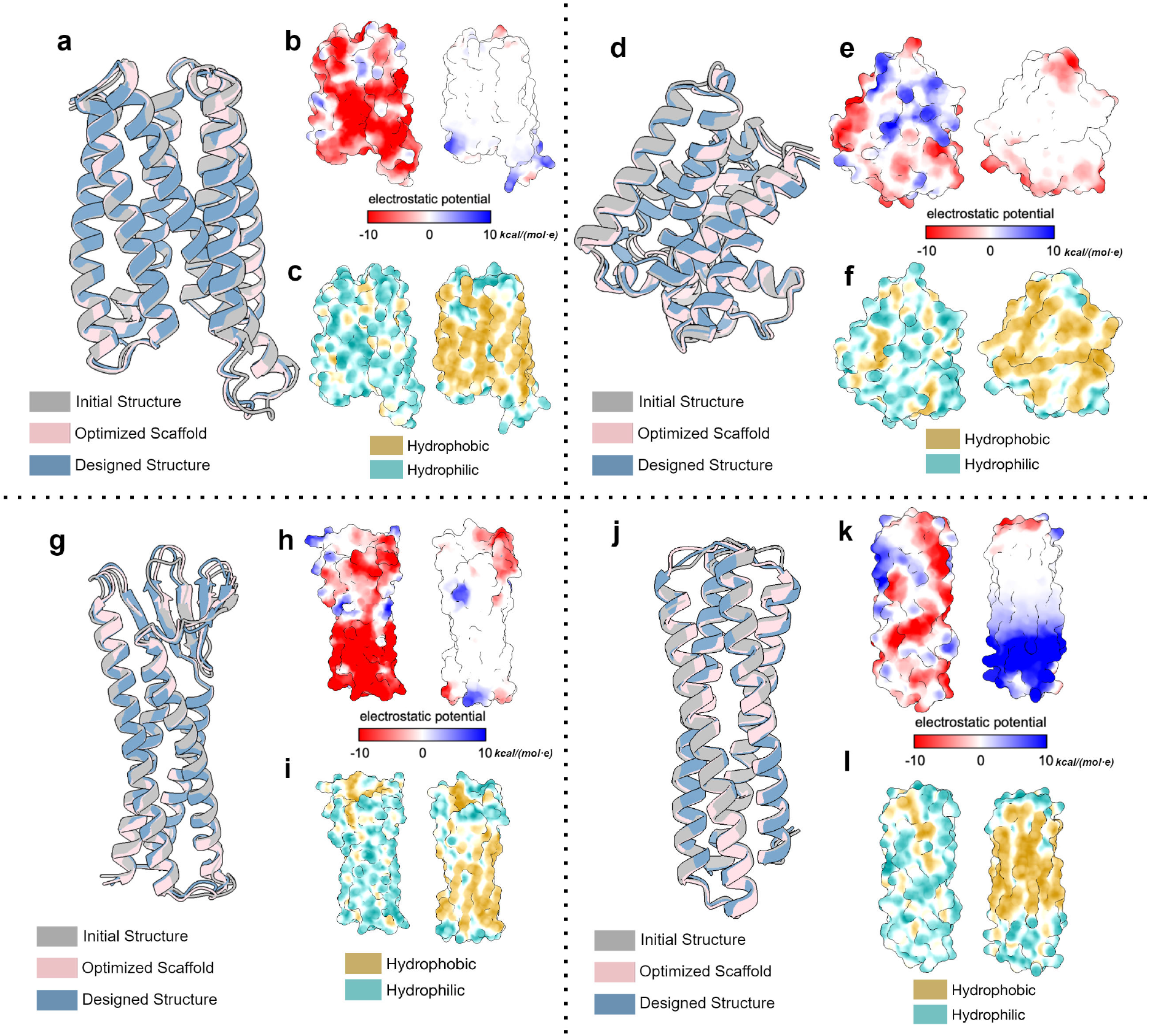
Visualization comparison of converted protein and initial protein. **a** Superimposition of the initial structure, optimized scaffold, and designed structure for 8oyy case. **b** Surface electrostatic potential before (initial) and after (designed) conversion for 8oyy (values in kcal/(mol · e) at 298 K). **c** Surface hydrophobicity before and after conversion for 8oyy. **d-l**, similar to a-c, but for the 8oyw, 8oyv, and 8w6f cases, respectively.

To assess structural consistency, we calculate the scRMSD of the designed structure with respect to the optimized scaffold (scRMSD_scaffold_), as detailed in the Methods section. The initial structure undergoes iterative processing through the pipeline to yield the optimized scaffold, which is then subjected to sequence redesign using MemProtMPNN; the ESMFold-predicted structure from this redesigned sequence is termed the designed structure. The scRMSD_scaffold_ is 0.506 Å, indicating a high design quality of the redesigned sequence. Meanwhile, based on the ESMFold prediction results, the protein’s pLDDT is 0.935, indicating a high level of confidence in our designed sequence.

### 2.3.2 Membrane insertion of the four cases

We further assessed the membrane insertion potential of the four cases using PPM 2.0. ^26^ As shown in Fig. 4a, after the MemConverter pipeline, the soluble GPCR-like protein (PDB: 8oyy) favorably integrates into the membrane plane, as demonstrated by a free energy change of −96.5 kcal/mol. Similar results were obtained for the other three cases, as illustrated in Fig. 4b-d and Table 1. Additionally, we utilized ProtRAP-LM to predict the membrane contact probability (MCP) of the protein sequence post-conversion (Fig. 4), finding excellent agreement with the predictions from PPM 2.0. Residues at the membrane surface consistently exhibited high MCP values.

**Table 1:**
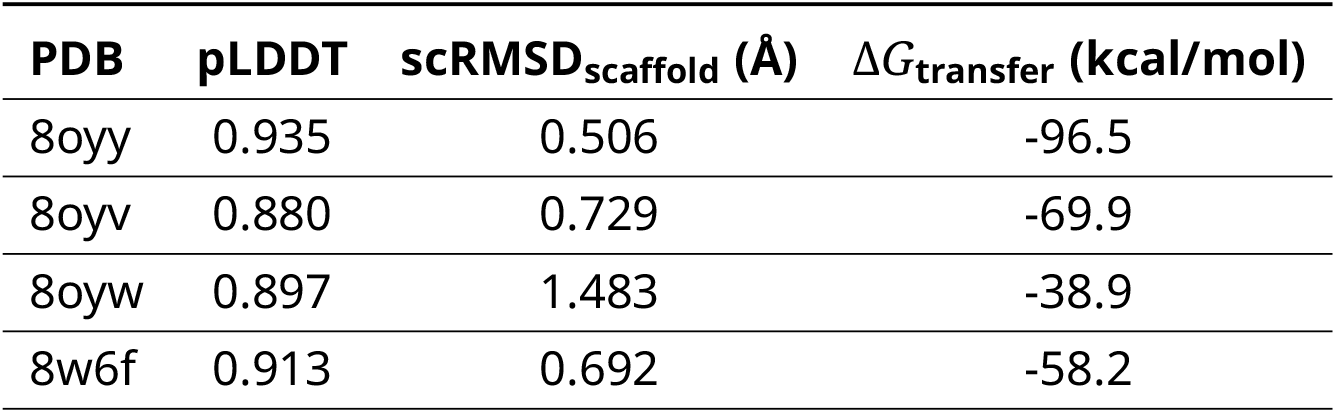
Quantitative Metrics for Protein Conversion Analysis.

**Figure 4:**
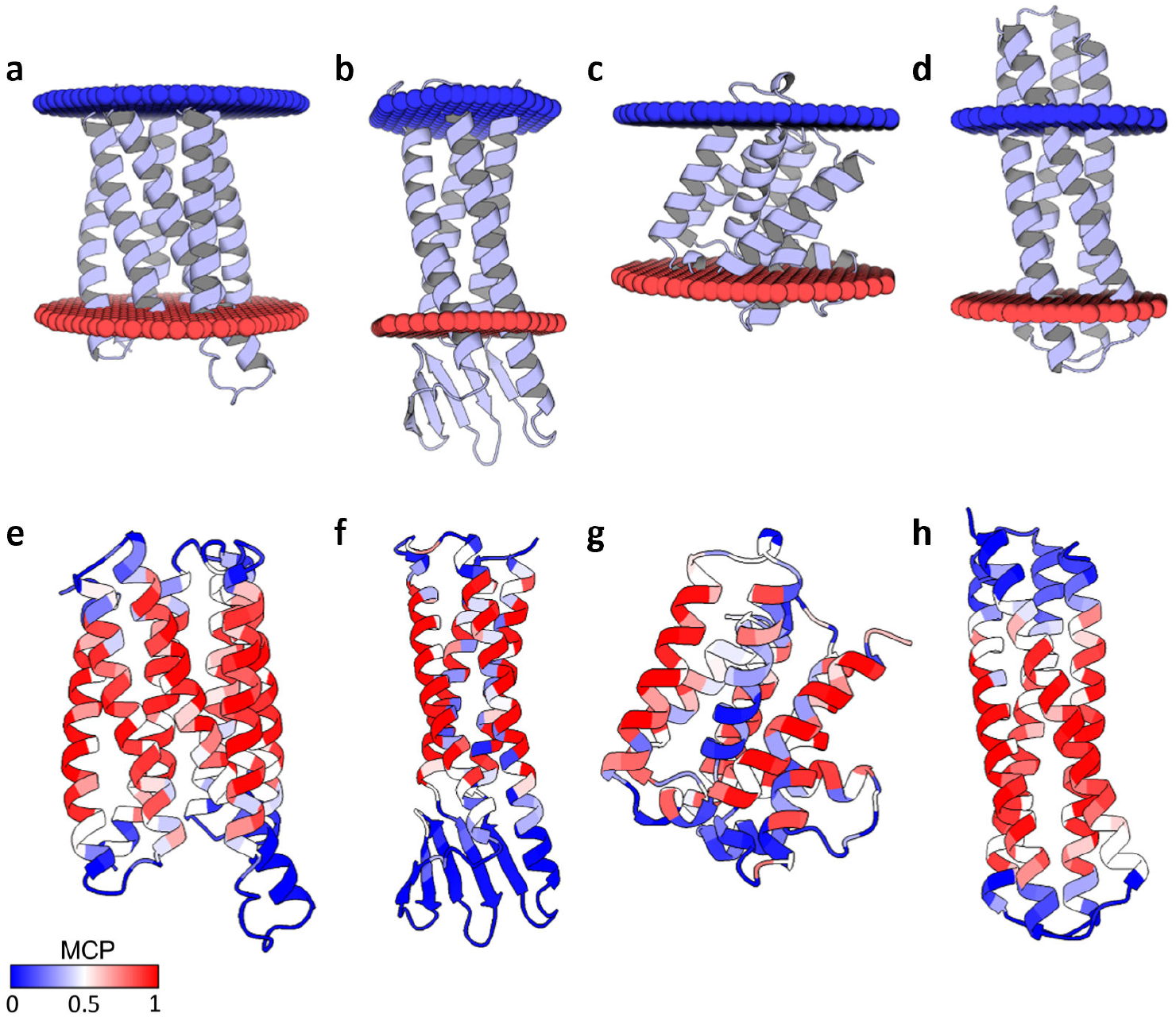
Predicted membrane interactions of the converted protein. **a**-**d** Localization of the designed proteins in the membrane as predicted by PPM 2.0, for the 8oyy, 8oyw, 8oyv, and 8w6f cases, respectively. **e**-**h** Membrane contact probabilities as predicted by ProtRAP-LM for the same cases.

To further validate the membrane localization of the designed proteins, we conducted *in silico* experiments using coarse-grained self-assembly MD simulations (Fig. 5). Initially, lipid and water molecules were randomly placed around the studied proteins (Fig. 5, left panels). When the initial proteins were placed in the simulation box, after the formation of the self-assembled lipid bilayer, the proteins will stay in the aqueous solution (Fig. 5, middle panels). In contrast, the designed proteins quickly integrated into the lipid bilayer membranes in 11 out of 12 replicate simulations (Fig. 5, right panels).

**Figure 5:**
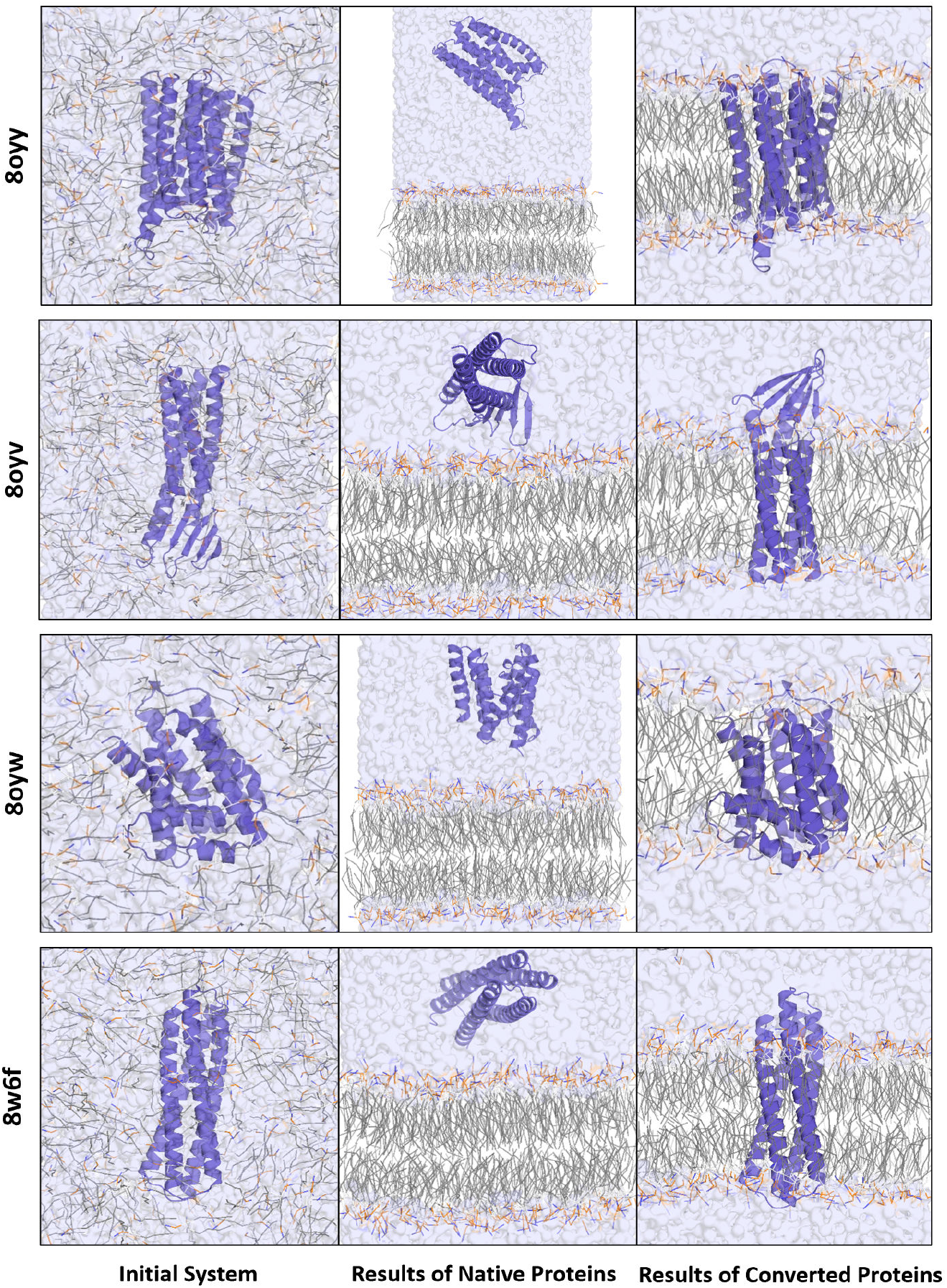
Validation of *soluble-to-membrane* conversion using coarse-grained self-assembly molecular dynamics simulations. The simulations compare the final localization of the native (initial) proteins and the converted (designed) proteins. The left column shows the initial state with proteins and lipids randomly distributed. The middle column shows that after simulation, the native proteins remain in the aqueous solution, failing to integrate with the self-assembled lipid bilayer. In contrast, the right column demonstrates that the converted proteins spontaneously insert and stabilize within the lipid bilayer. This in-silico validation supports the successful conversion from soluble proteins to membrane proteins.

All four studied cases exhibited similar results, as detailed in Table 1. The designed proteins display high-quality sequences, significant structural self-consistency, and low transfer free energy as detected by PPM 2.0. These findings underscore the robustness of our method and its applicability to a variety of systems.

#### 2.3.3 Performance evaluation

To quantitatively evaluate the effectiveness of our method and the advantages of using MemConverter for protein design, we conducted systematic tests for the four cases described above.

Four sequence design methods were evaluated. 1) As a baseline, we first employed ProteinMPNN to directly design sequences for the initial structure, without utilizing the MemConverter pipeline. 2) Designing sequences for the initial structure using MemProtMPNN, without utilizing the MemConverter pipeline. 3) Designing sequences using the MemConverter pipeline, with ProteinMPNN employed for sequence generation. 4) Designing sequences using the MemConverter pipeline, with MemProtMPNN utilized for sequence generation. With each method, we generated 36 sequences for each of the four proteins and predicted their structures using ESMFold. We then extracted pLDDT, calculated the RMSD to the optimized scaffold (scRMSD_scaffold_), and finally used PPM 2.0 to predict the membrane insertion free energy. The results are presented in Fig. 6.

**Figure 6:**
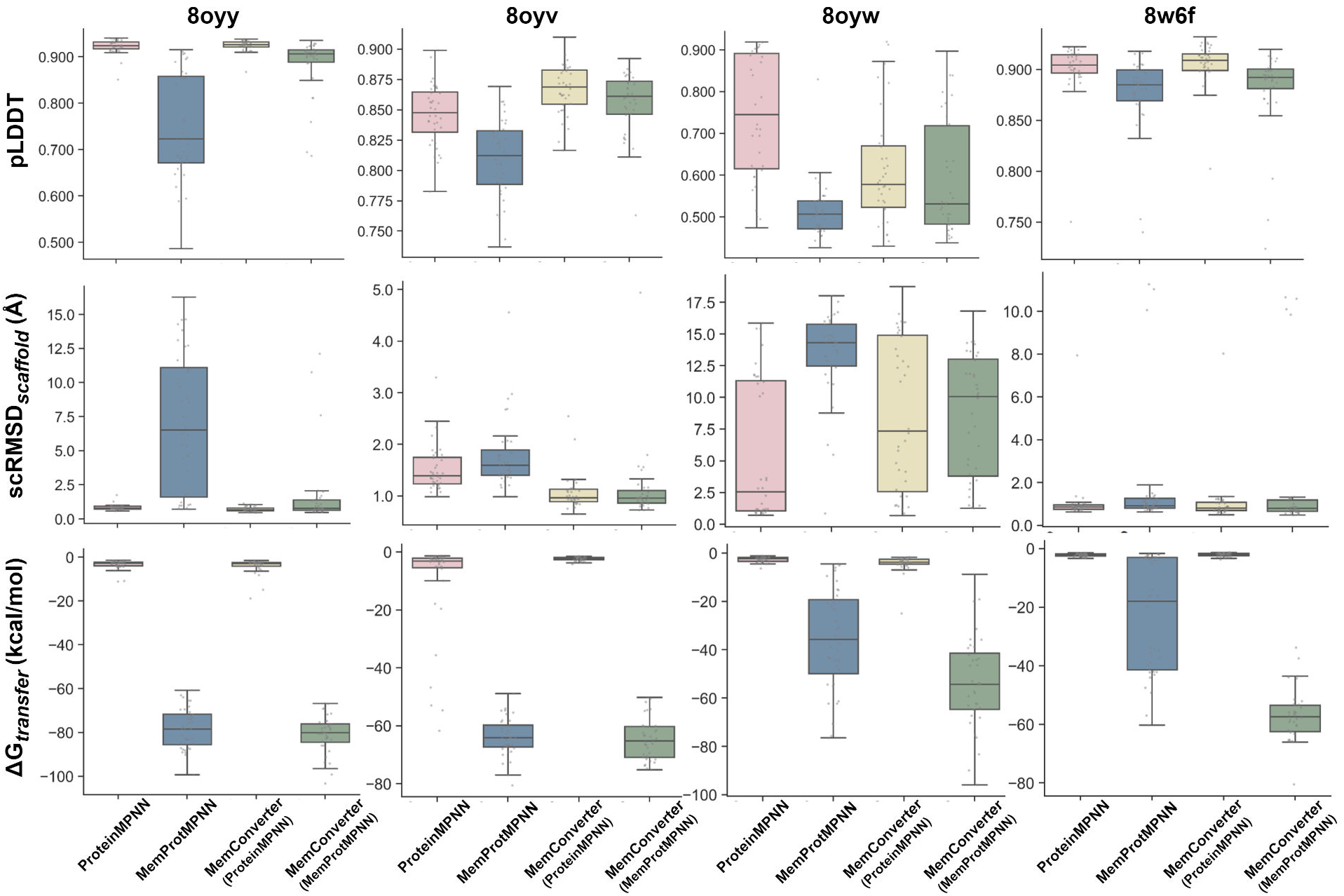
Quantitative comparison of four different methods for converting soluble proteins to membrane proteins across four case studies (PDB IDs: 8oyy, 8oyv, 8oyw, 8w6f). For each case and method, 36 sequences were generated and evaluated via ESMFold-predicted structures and PPM 2.0. Panels show distributions of pLDDT, scRMSD_scaffold_, and PPM 2.0 transfer free energy Δ*G*_transfer_. The comparison reveals that methods using ProteinMPNN (Method 1,3) yield high structural quality (high pLDDT, low *scRMSD*_*scaffold*_) but fail to achieve membrane insertion (Δ*G*_*transfer*_ values near zero). Conversely, applying MemProtMPNN directly to the initial structure (Method 2) achieves favorable insertion but suffers from poor structural quality (low pLDDT, high *scRMSD*_*scaffold*_). The results demonstrate that only the complete MemConverter(MemProtMPNN) pipeline (Method 4) successfully achieves both high structural quality and favorable membrane insertion simultaneously. Box plots indicate medians, quartiles, and outliers.

Comparative analysis in Fig. 6 reveals that sequences designed by ProteinMPNN, while potentially of higher quality, are typically soluble protein sequences. As confirmed by the PPM analysis, these sequences do not tend to be integrated into the membrane. Conversely, designing sequences with the initial protein backbone directly using MemProtMPNN yields membrane protein sequences that, while successfully integrating into the membrane as detected by PPM 2.0, exhibit lower pLDDT scores and higher scRMSD_scaffold_ values, suggesting compromised design quality and low structural consistency. In contrast, employing the MemConverter pipeline with MemProtMPNN to design sequences not only achieves higher structural quality and structural consistency but also allows PPM 2.0 to identify them as membrane proteins.

Specifically, for the 8oyy case, directly designing sequences for the initial structure or optimized scaffold using ProteinMPNN typically yields high average pLDDT values of 0.923 and 0.924, respectively, with scRMSD values within 1 Å, achieving atomic-level precision. However, the average membrane transfer free energy is approximately −3 kcal/mol. This indicates that proteins designed directly with ProteinMPNN are unlikely to spontaneously integrate into the membrane. We speculate that this is due to ProteinMPNN’s tendency to design soluble proteins, as the majority of its training set consists of soluble proteins. Consequently, while the designed sequences are of high structural quality, they do not achieve the highly negative transfer free energy required for membrane insertion.

In contrast, designing sequences for the initial structure using MemProtMPNN results in an average membrane transfer free energy of approximately −78 kcal/mol, accompanied by a lower average pLDDT of 0.723 and a standard deviation of 0.114. This indicates that applying MemProtMPNN directly to the initial structure, which has not yet been optimized for a membrane environment, leads to lower-quality designs. Encouragingly, when the MemConverter pipeline combined with MemProtMPNN is employed to design sequences, a structure better adapted to the membrane environment is generated during the iteration. As a result, we were able to achieve both high design quality (average pLDDT of 0.907, scRMSD_scaffold_ of 0.770) and a favorable spontaneous transfer free energy of approximately −80 kcal/mol.

### 2.4 Conversion of membrane proteins to soluble proteins

Through subtle adjustments, we can achieve the reverse transformation—from membrane proteins to soluble proteins. Specifically, we set the target MCP to 0 for all the residues and replace MemProtMPNN with SolubleMPNN as the sequence generator. To demonstrate this reverse conversion, we conducted experiments on several membrane proteins, including GPCR (PDB: 6ffi) and Claudin (PDB: 4p79).

Apart from the aforementioned adjustment, the MemConverter pipeline remains fundamentally unchanged, wherein we design a series of sequences and select high-quality candidates based on pLDDT scores. Fig. 7a illustrates significant surface remodeling during the conversion, characterized by a transition from hydrophobic to hydrophilic surfaces and an increase in polar or charged residues. Furthermore, based on the ESMFold prediction results, the protein’s pLDDT is above 0.91, indicating a high level of confidence in the designed sequences.

**Figure 7:**
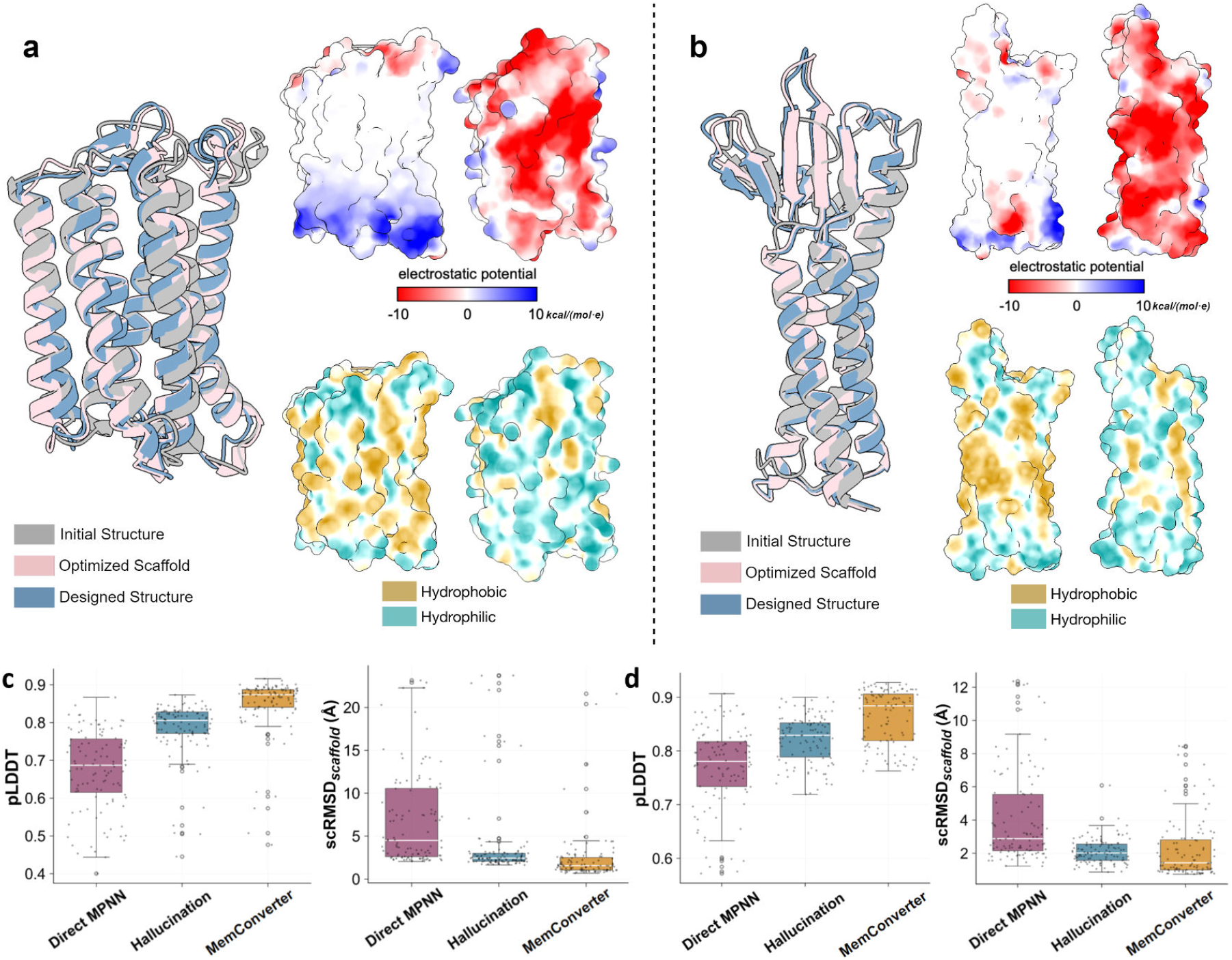
**a**,**b** Visualization of membrane-soluble transformation results for the structures with PDB IDs 6ffi (**a**) and 4p79 (**b**). Superimposed of initial structure, optimized scaffold, and designed structure are shown alongside surface electrostatic potential (values in kcal/(mol · e) at 298 K) and hydrophobicity. **c**,**d** Quantitative comparison of conversion methods (Direct, Hallucination, and MemConverter) for the 6ffi and 4p79 cases.

To evaluate the effectiveness of our method, we compared three design approaches. (1) Direct sequence design utilizing SolubleMPNN. (2) Hallucination-optimized structures paired with SolubleMPNN, as described by Goverde et al. ^20^ (3) Our MemConverter pipeline in combination with SolubleMPNN. For method 1, we generated 108 sequences for the initial membrane protein using SolubleMPNN and evaluated their pLDDT scores with ESMFold. For the latter two methods, we first designed three backbones for each initial protein, then generated 36 sequences per backbone and assessed sequence quality and the potential for successful folding using ESMFold.

Analysis of pLDDT score distributions reveals that MemConverter yields structures of superior quality, as shown in Fig. 7c and Fig. 7d. Notably, MemConverter generates a scaffold in 1–2 minutes using a single NVIDIA A40 GPU, requiring only a few AlphaFold2 inference passes. In contrast, hallucination-based backbone generation typically demands hundreds of inference passes and backpropagation, consuming 15–20 minutes.

We further validated the solubility of the designed proteins through coarse-grained self-assembly MD simulations. Similar to the previous section, the studied proteins, lipids, and water molecules were initially randomly placed in a simulation box (Fig. 8, left panels). After 100 ns of coarse-grained MD simulations, the initial proteins integrated stably into the membrane through self-assembly (Fig. 8, middle panels), whereas the converted proteins were found in the aqueous solution (Fig. 8, right panels). This *in-silico* validation supports the successful conversion from transmembrane proteins to soluble proteins.

**Figure 8:**
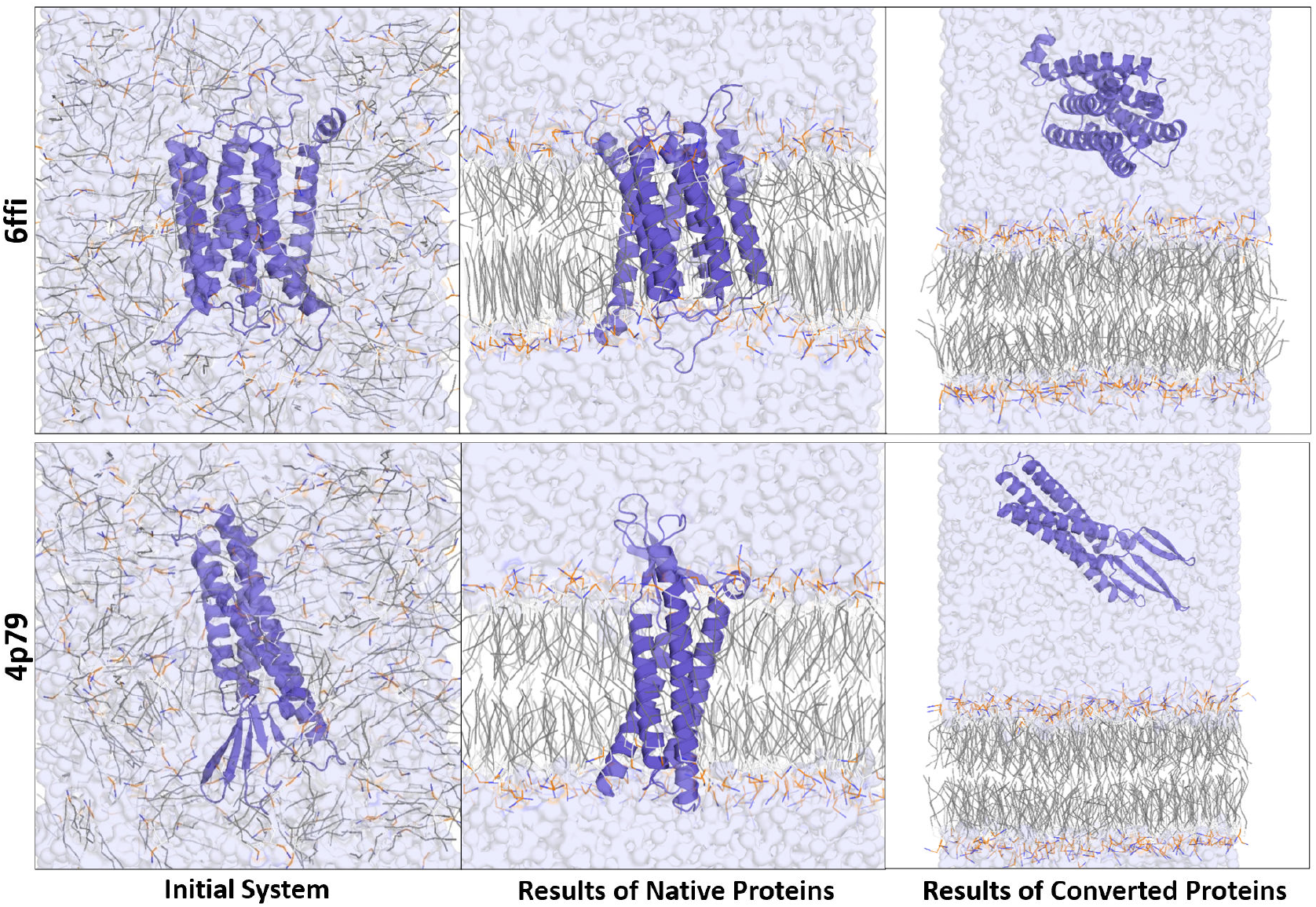
Validation of *membrane-to-soluble* conversion using coarse-grained self-assembly molecular dynamics simulations. The simulations compare the final localization of the native membrane proteins and their converted soluble analogues. The left column shows the initial state with proteins and lipids randomly distributed. The middle column shows that after simulation, the native proteins integrated stably into the self-assembled lipid bilayer. In contrast, the right column demonstrates that the converted proteins remain in the aqueous solution and do not integrate into the membrane. This in-silico validation supports the successful conversion from transmembrane proteins to soluble proteins.

These results underscore the versatility and robustness of the MemConverter pipeline as a universal tool for reprogramming the surface properties of proteins for targeted localization. The pipeline is applicable to both transmembrane and soluble proteins.

## 3 Discussion

Although our pipeline has not yet undergone experimental validation, the structure predictions and molecular dynamics simulations provide compelling evidence for its eficacy in generating stable membrane proteins and achieving successful bidirectional conversions between soluble and membrane proteins. The designed proteins fulfill our design objectives and demonstrate stable integration within lipid bilayers or exhibit enhanced solubility. These results highlight the potential of our MCP-guided MemConverter pipeline to precisely manipulate protein surface properties for the desired localization.

The pipeline’s bidirectional capability, enabling both membrane-to-soluble and soluble-to-membrane conversions, provides a versatile framework for addressing various protein engineering challenges. This flexibility facilitates applications such as the design of soluble analogs of membrane proteins for structural studies, as well as the engineering of membrane proteins with tailored functionalities for therapeutic purposes.

A key strength of our pipeline lies in its high extensibility. The modular architecture allows for the seamless replacement of the sequence design model, MemProtMPNN, with other membrane protein design tools as they become available. This adaptability ensures that the pipeline can evolve alongside advancements in computational protein design, thereby enhancing its capacity to generate novel membrane proteins with improved accuracy and efficiency.

However, challenges remain, including the need for systematic experimental validation to confirm the functionality and stability of the designed proteins *in vitro* and *in vivo*. Future work could also explore the incorporation of additional environmental constraints, such as specific lipid compositions or pH conditions, to enhance the biological applicability of the designs.

Notably, while our pipeline yields competitive metrics for overall structural consistency between the final designed protein and the initial scaffold (scRMSD_initial_), it does not invariably achieve the lowest values among the compared methods (Fig. 9). We posit that this reflects a necessary compromise inherent to adapting soluble proteins for membrane environments. Indeed, omitting structural iteration and directly applying MemProtMPNN to redesign soluble proteins results in a substantial decline in the quality of generated sequences. As discussed in the Introduction section, conversions between membrane and soluble proteins often necessitate a degree of structural divergence to accommodate environmental demands. Accordingly, our pipeline seeks an optimal balance between these constraints, enabling designed proteins to adapt to the target localization’s topological requirements while preserving the core architecture of the initial scaffold to the greatest extent feasible. Nonetheless, the precise impact of membrane environments on such structural perturbations warrants further investigation.

**Figure 9:**
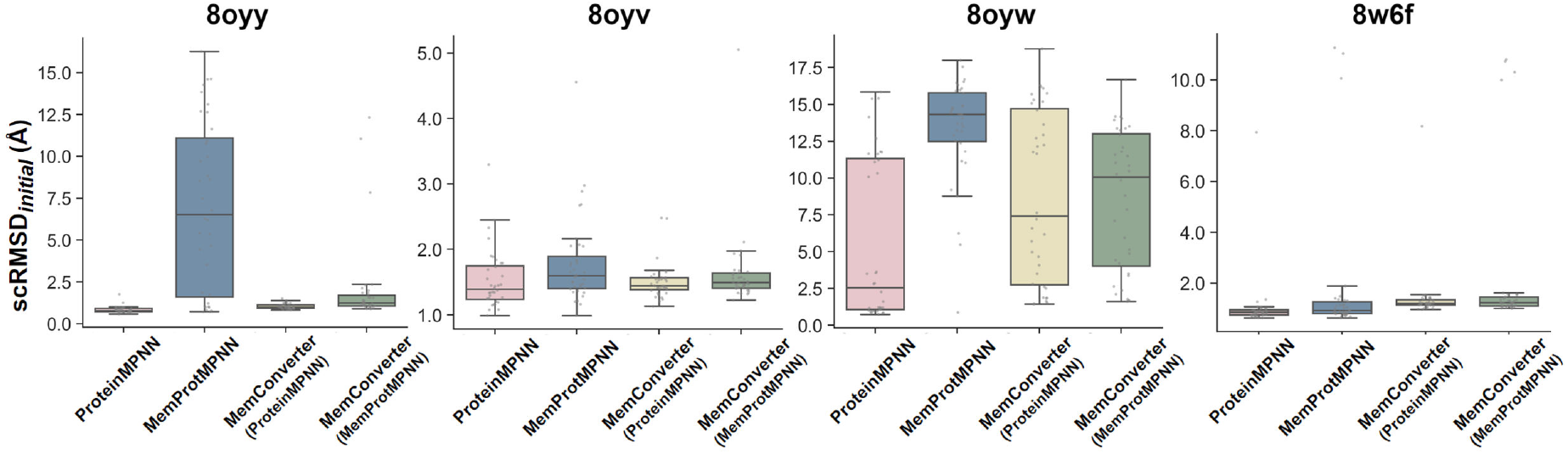
scRMSD_initial_ of four different methods for converting soluble proteins to membrane proteins across four case studies (PDB IDs: 8oyy, 8oyv, 8oyw, 8w6f).

Despite these constraints, our pipeline represents a robust and adaptable tool for protein design, with the potential to significantly advance the field of membrane protein engineering. Its modular design and reliance on MCP-guided sequence optimization position it as a forward-looking framework, capable of leveraging future computational innovations to address complex protein design challenges.

## 4 Methods

### 4.1 Datasets, Training, and Evaluation for MemProtMPNN

#### Datasets Construction

We fine-tuned this model using a curated dataset of membrane proteins from the transmembrane AlphaFold Database (tmAFDB). ^21^ A key chal-

This metric measures *C*_*α*_ RMSD relative to the initial structure, assessing core topology preservation. The plots demonstrate that while directly applying MemProtMPNN to the initial structure leads to significant structural divergence (high *scRMSD*_*initial*_), the MemConverter pipeline successfully preserves the core architecture. The slightly higher *scRMSD*_*initial*_ for the pipeline using MemProtMPNN compared to Protein-MPNN reflects a necessary structural compromise, enabling the protein to adapt to the topological requirements of the membrane environment.

lenge is that, for many proteins, transmembrane segments constitute only a fraction of the overall structures, yet they possess markedly distinct topologies compared to soluble domains. Therefore, to specialize the model for the unique topological constraints of membrane proteins, our training was performed exclusively on isolated transmembrane domains extracted from tmAFDB.

To construct a dataset of transmembrane domains, the domain annotations from The Encyclopedia of Domains (TED) ^27^ were employed. For proteins lacking TED annotations, Merizo^28^ was used to identify domains. The dataset was subsequently filtered using TMbed^25^ to exclude proteins with fewer than 20% transmembrane residues or domains shorter than 32 residues. This curation process resulted in a final dataset comprising 85,051 domains for training, validation, and testing. The length distribution of the dataset is shown in Fig. S1.

The curated dataset of 85,051 membrane protein domains was divided into training (85%, 72,293 domains), validation (9%, 7,655 domains), and test (6%, 5,103 domains) sets. MMseqs2^29^ was employed for clustering at 30% sequence similarity (coverage: 0.8, cov_mode: 0). Proteins within the same cluster were assigned exclusively to one dataset partition to prevent data leakage.

#### Model Training

MemProtMPNN training followed the protocol established for HyperMPNN, ^30^ utilizing ProteinMPNN architecture for inverse folding (https://github.com/meilerlab/HyperMPNN). Training incorporated 0.2 Å Gaussian noise on backbone coordinates, 10% dropout, a batch size of 3000, 500 epochs and 1000 examples per epoch.

#### Model Evaluation

MemProtMPNN was evaluated on a simplified test set comprising 3,722 transmembrane proteins, with one representative sequence retained per cluster to ensure diversity and avoid redundancy. The metrics employed are as follows.

pLDDT and scRMSD are commonly employed metrics for evaluating the designability of sequence generation models. ^7,31,32^ These scores are derived by generating a single sequence using either MemProtMPNN or ProteinMPNN, followed by structure prediction with ESMFold ^33^ to compute the corresponding pLDDT value, along with the C*α*-RMSD relative to the initial structure.

Sequence recovery serves as a measure of the fidelity with which a model recapitulates the native amino acid sequence when provided with a fixed protein backbone. ^7^ It is formally defined as the percentage of native residues correctly recovered in the model’s predicted sequence relative to the total number of residues in the test set.

Amino acid frequency deviations (also termed amino acid compositional bias) quantify the discrepancies in amino acid composition between designed sequences and corresponding frequencies observed in native PDB structures, thereby assessing potential biases in amino acid preferences. ^7^ Specifically, these deviations are calculated as the differences between the observed frequency of each amino acid in the designed sequences and its average frequency in the native sequences.

Diversity quantifies the extent of sequence variability elicited by the model for a given fixed backbone, thereby serving as a proxy for the accessible volume of the compatible sequence space. ^7^ For each backbone, *N* = 32 sequences are sampled at a temperature of 0.5; the pairwise sequence identity (fraction of identical residues) is

subsequently computed for all sequence pairs, yielding a per-protein diversity score of 1 − *Ī*, where *Ī* represents the mean pairwise identity. The global diversity is then reported as the mean of these scores across the test set.

### 4.2 Pipeline Implementation

Our design methodology was implemented based on the AF2cycler framework,^23^ as implemented in ColabDesign.

#### Sequence Generation with MemProtMPNN

During the iterative design process, protein sequences were represented as logits. For a given protein backbone of length *L*, a tensor of shape (*L*, 20) was used to represent the score for one of the 20 canonical amino acids at each position. The sequences designed by either MemProtMPNN or SolubleMPNN were propagated to the subsequent cycle in the logit format.

#### Residue Freezing

To ensure that the design effort can be concentrated on membrane-interacting regions while allowing other residues to be fixed during the iterative pro-cesses (Fig. 10c), certain residues were constrained according to two criteria: membrane-relative spatial constraints and solvent accessibility. To standardize protein orientation for spatial filtering, we first aligned the primary principal axis of the protein with the Z-axis using a Principal Component Analysis (PCA)-based method adapted from PyMOL’s orient function^34^ (see Algorithm 1 in the Supplementary Information). A hypothetical membrane plane was then defined by user-specified boundaries along the Z-axis, and all residues outside this region were frozen. Concurrently, residues predicted to be buried within the protein’s hydrophobic core were also fixed. Buried residues were identified using Relative Accessible Surface Area (RASA) values predicted by the ProtRAP-LM (https://github.com/ComputBiophys/ProtRAP-LM); ^19^ residues with a RASA score below 0.35 were considered buried and remained unchanged throughout the design process.

**Figure 10:**
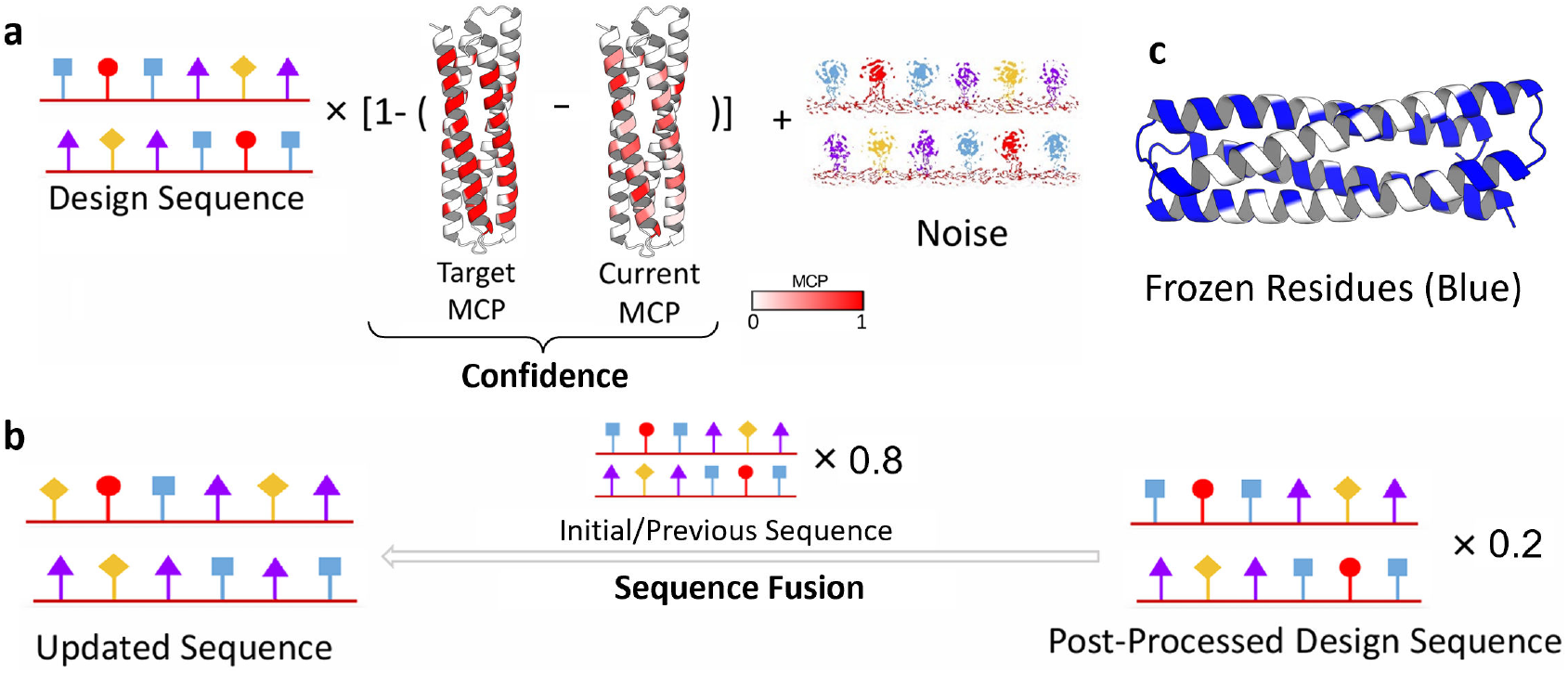
MCP-Guided logits fusion. **a** MCP-guided sequence logits processing. The logits of the designed sequence are perturbed based on the MCP value to ensure that the sequence evolves in the desired direction. **b** Sequence logits fusion. The logits of the post-processed design sequence are combined with the logits from the previous iteration in a weighted manner to ensure gradual evolution. **c** Frozen residues. Residues in non-transmembrane regions or buried residues with low RASA are fixed from mutations during iterations (shown in blue).

#### MCP-Guided Confidence Calculation

Before initiating the iterations, the target MCP of each residue is established for the protein based on the user-defined transmembrane region. Then, the MCP of the current sequence is predicted using ProtRAP-LM. ^19^ The confidence for each residue is determined by the discrepancy between the predicted MCP and the target MCP, as calculated by 1 − | *MCP*_*target*_− *MCP*_*current*_ |.

For the conversion of a membrane protein to a soluble protein, the target MCP is 0. The confidence score (*𝒞*) was therefore calculated as:

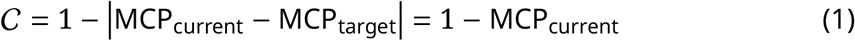

For the conversion of a soluble protein to a membrane protein, where RASA values are typically close to 1, the confidence score was approximated using the Relative Lipid Accessibility (RLA) for indirect guidance. RLA, which quantifies the fraction of a residue’s accessible surface area exposed to lipid environments, is computed as RLA = RASA × MCP. For unfreeze residues, MCP_target_ = 1, and thus the C for these residues can be written as the following expression.

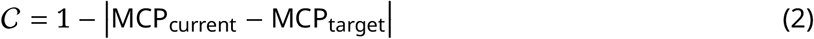

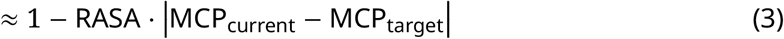

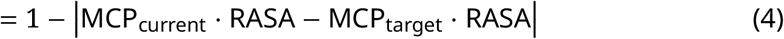

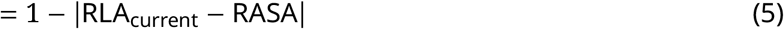

All biophysical properties (RASA, RLA, MCP) were predicted using ProtRAP-LM. ^19^

#### Sequence Fusion

To prevent structural collapse due to significant changes in the sequence, it is not advisable to use the new designed sequence as the starting point for the subsequent iteration. Instead, we combined them with the logits from the previous (or initial) sequence using a weighted fusion (e.g., a weight of 0.2 for the new logits and 0.8 for the previous logits) (Fig. 10b). This step ensures that the sequence evolves gradually.

#### Iterative Structure Prediction

Structure prediction within each design cycle was performed using structure prediction model in ColabDesign https://github.com/sokrypton/ColabDesign. To ensure structural consistency and accelerate convergence, the structure from the preceding iteration was employed as both a structural template (`use_templates=True`) and the initial atomic coordinates (`use_initial_atom_pos=True`). For proteins with substantial extramembrane domains, the initial input structure was selectively utilized as a template for these regions to preserve their native conformation.

#### Monitoring of Transmembrane Signal

The emergence of transmembrane features during the design trajectory was monitored using TMBed(https://github.com/BernhoferM/TMbed), a prediction tool independent of ProtRAP-LM. The iterative process was terminated once the designed sequences satisfied a predefined criterion: over 80% of residues in the user-defined membrane plane were detected as transmembrane residues by TMBed.

#### Assessment of Sequence Quality and Structural Consistency

During the screening process, the quality of the final designed sequences was evaluated using multiple metrics. First, the structure of designed sequences was predicted with ESMFold(https://github.com/facebookresearch/esm). The folding confidence was assessed by the average pLDDT of C*α* atoms from the ESMFold prediction. Second, structural consistency was quantified by the C*α* -Root Mean Square Deviation (C*α* -RMSD) between the ESMFold-predicted structure and the initial input structure, calculated using the `Superimposer` module from the `Bio.PDB` ^35^ library.

To evaluate the structural consistency of the redesigned sequences for the optimized scaffold, scRMSD_scaffold_ is employed, which quantifies the C*α* root-mean-square deviation (RMSD) between the iteratively optimized scaffold and the ESMFold-predicted structure of the designed sequence, with lower values indicating faithful folding to the target backbone. In the discussion section, we employed another metric: scRMSD_initial_, which measures the C*α* RMSD between the initial structure and the designed structure. Both metrics are illustrated in Fig. S2.

#### Assessment of Transmembrane Propensity

The propensity of a designed protein to spontaneously integrate into a lipid bilayer was quantified by its transfer free energy (Δ*G*_transfer_). This value, representing the free energy change from an aqueous to a membrane environment, was calculated using a local version of the PPM 2.0 server(https://cggit.cc.lehigh.edu/biomembhub/PPM2.0_server_code). The PPM method determines the optimal protein orientation within the membrane by finding the position that minimizes the transfer free energy. A Δ*G*_transfer_ value in the range of −10 to −400 kcal/mol is typically characteristic of a stable integral membrane protein.

### 4.3 Coarse-grained Self-assembly Simulation

Molecular dynamics simulations were performed using the GROMACS 2023.4 package, ^36^ employing the Martini 3 coarse-grained force field.

System preparation began with the initial protein all-atom structures. For proteins with missing structural elements, such as loops or terminal residues, the models were first completed using the PDBFixer tool from the OpenMM 7 library. ^37^ Following this preprocessing step, the complete all-atom structures were converted into a coarsegrained representation using the `martinize2` script from the Python `vermouth` package. ^38^ An elastic network was then applied to maintain the protein’s tertiary structure. Subsequently, the coarse-grained protein was placed in a simulation box containing randomly distributed dipalmitoylphosphatidylcholine (DPPC) molecules. The entire system was solvated with Martini water and neutralized by adding Na^+^ or Cl^−^ counter-ions, with all steps performed using GROMACS utilities. The final system was contained within a simulation box of approximately 9 × 9 × 12 nm^3^.

Before the production simulations, each system was energetically relaxed through a two-stage protocol consisting of 1000 steps of energy minimization, followed by 400 ps of equilibration under the NPT ensemble.

For each protein, three independent production simulations (replicates) were conducted for a duration of 80 ns each, using an integration time step of 20 fs. The temperature was maintained at a constant 300 K using the v-rescale thermostat^39^ with a time constant of *τ*_*T*_ = 1 ps. The pressure was controlled at 1 bar using the Par-rinello–Rahman barostat^40^ with a time constant of *τ*_*P*_ = 12 ps. The pressure coupling was isotropic for the first 40 ns and was subsequently switched to semi-isotropic for the remaining 40 ns of the simulation. A cutoff distance of 1.1 nm was employed for both van der Waals and electrostatic interactions, with long-range electrostatics treated via the reaction field method.^41^

Protein visualizations in this paper were generated using PyMOL ^34^ and UCSF ChimeraX. ^42,43^ ChimeraX was specifically employed to render the electrostatic potential and surface hydrophobicity.

## 4.4 Data Availability Statement

The source code for MemConverter and associated materials will be available at https://github.com/ComputBiophys/MemConverter after peer review. For each case study protein, the ten selected designed sequences and their corresponding metrics (e.g., pLDDT, *ΔG*_transfer_, scRMSD_scaffold_, and scRMSD_initial_) are provided in the Supplementary Information (Table S1-S6).

## 5 Acknowledgements

The authors appreciate the study by Christopher Frank et al., “Alphafold2 refinement improves designability of large de novo proteins”, which served as a major inspiration for this work. The authors express their gratitude to Zefeng Zhu for insightful discussions. This work was supported by the National Key R&D Program of China (2024YFA0916800 to C.S.) and the Science Fund for Innovative Research Groups of the National Natural Science Foundation of China (T2321001 to C.S.). Part of the molecular dynamics simulations were performed on the computing platform of the Center for Life Sciences at Peking University.

## Supplementary Information for

### 1 Supplementary Algorithm

#### 1.1 PCA-based Protein Orientation

##### Algorithm 1

PCA-based Protein Orientation

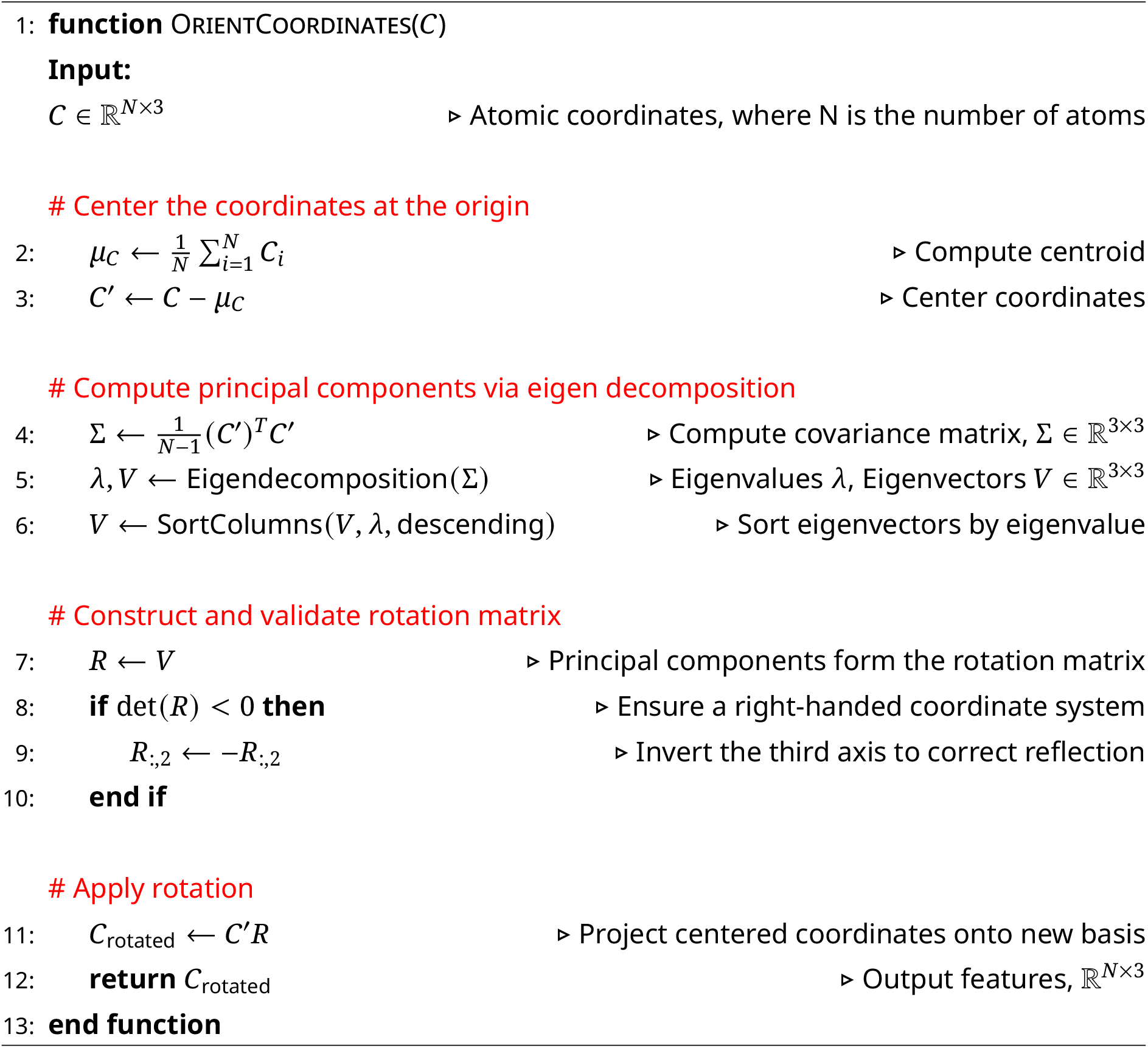

### 2 Supplementary Tables

**Table S1:**
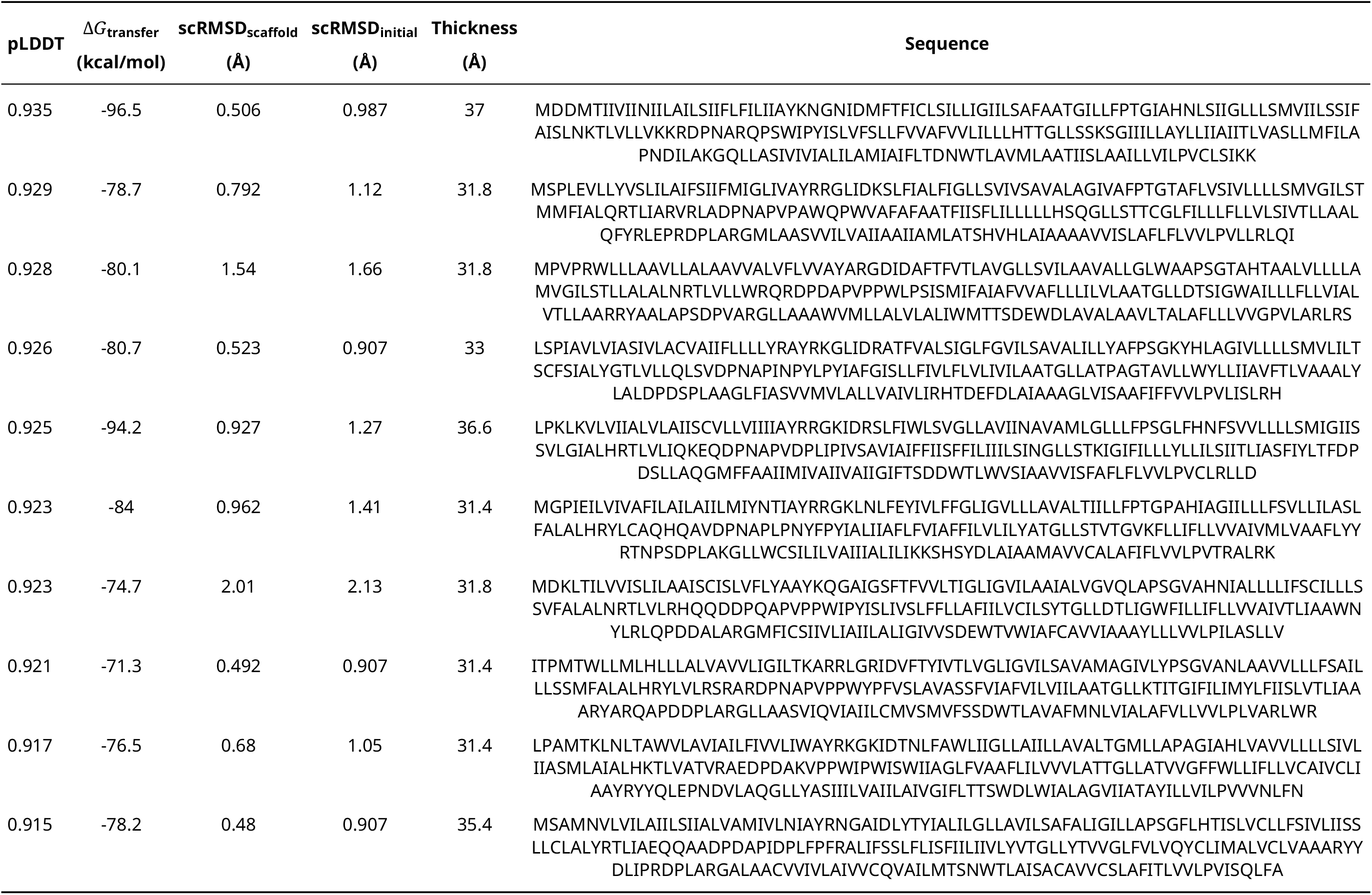
List of Design Sequences for Protein 8oyy’s Membrane Analogues.

**Table S2:**
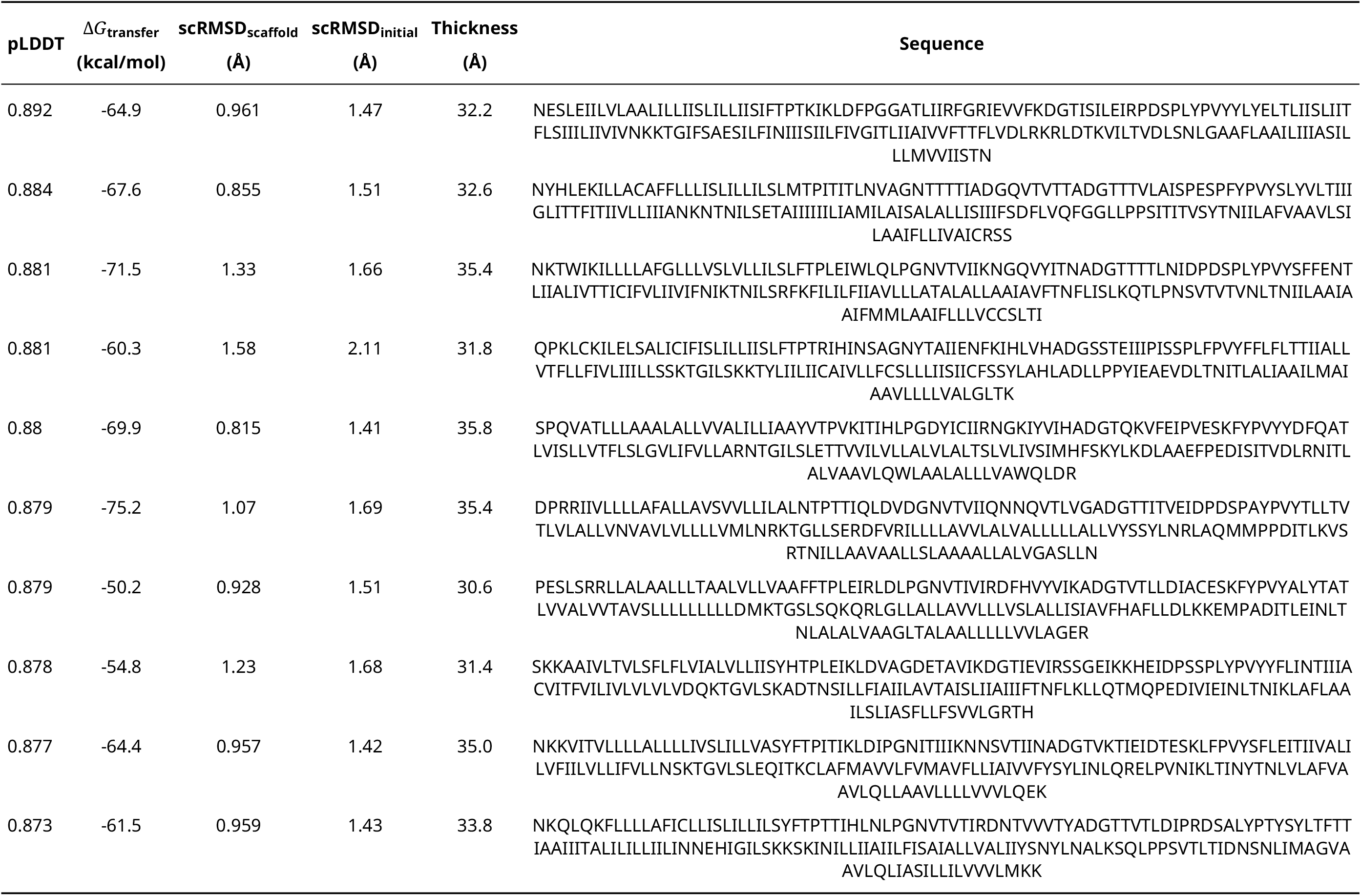
List of Design Sequences for Protein 8oyv’s Membrane Analogues.

**Table S3:**
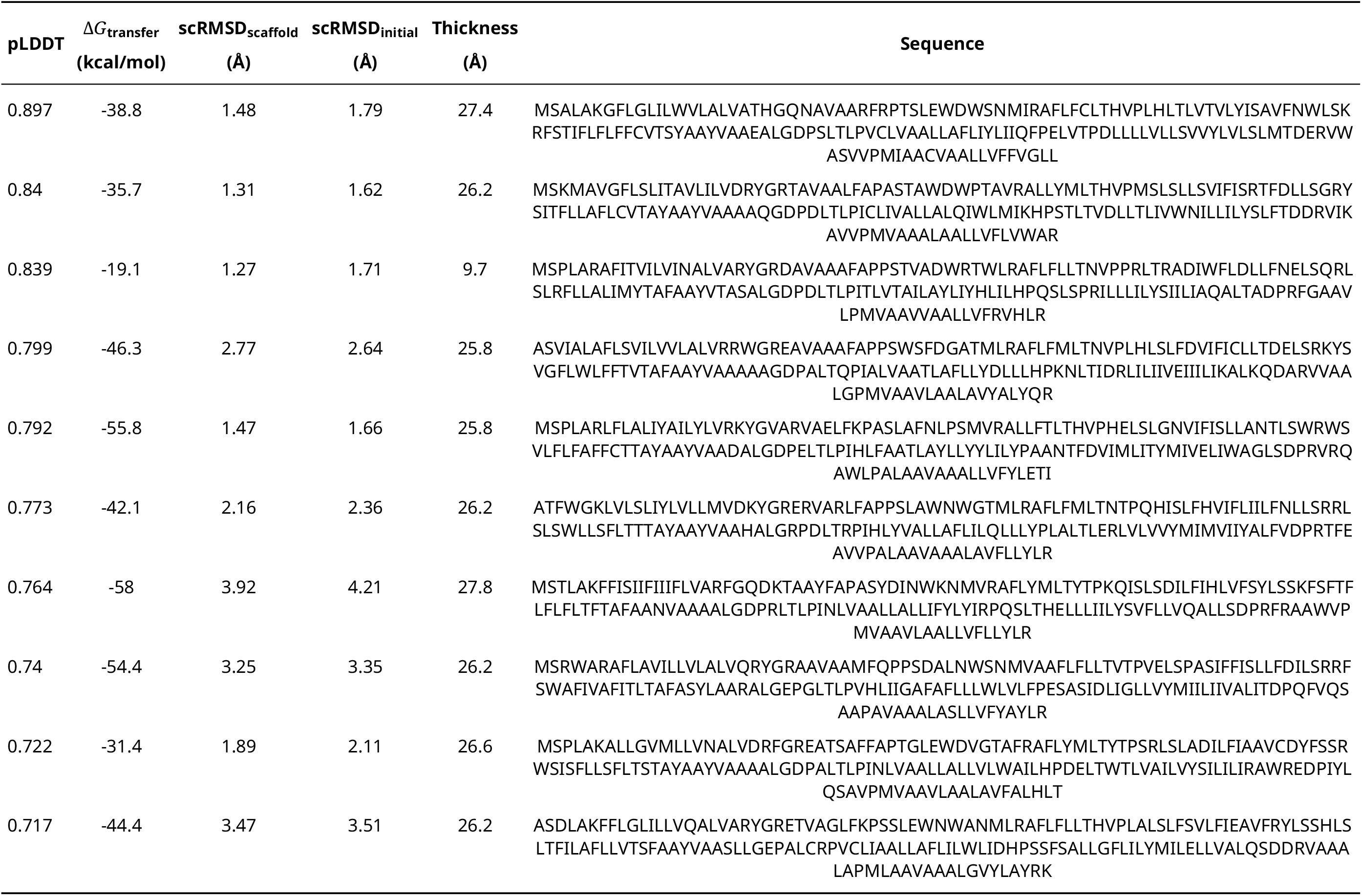
List of Design Sequences for Protein 8oyw’s Membrane Analogues.

**Table S4:**
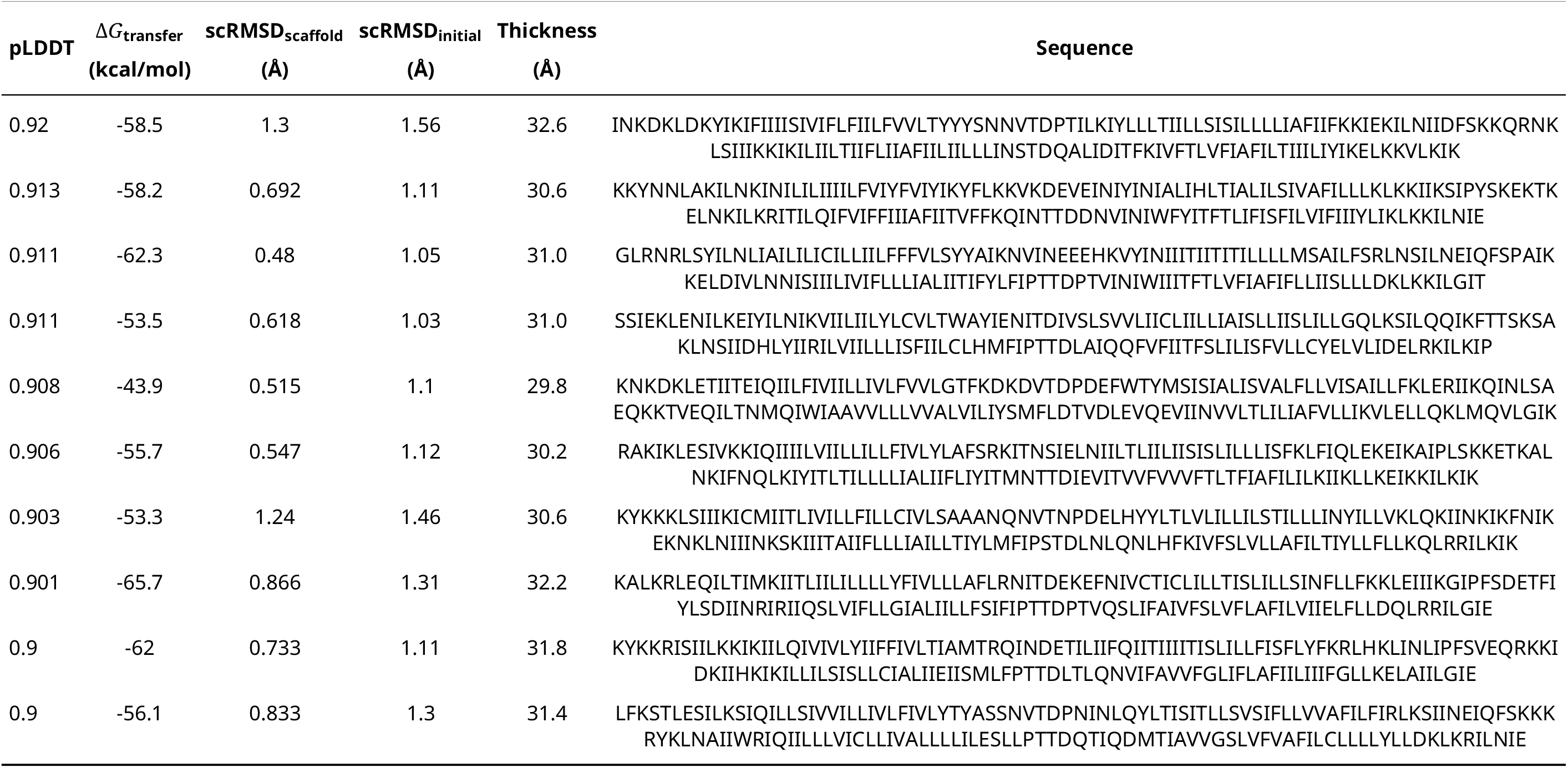
List of Design Sequences for Protein 8w6ffi’s Membrane Analogues.

**Table S5:**
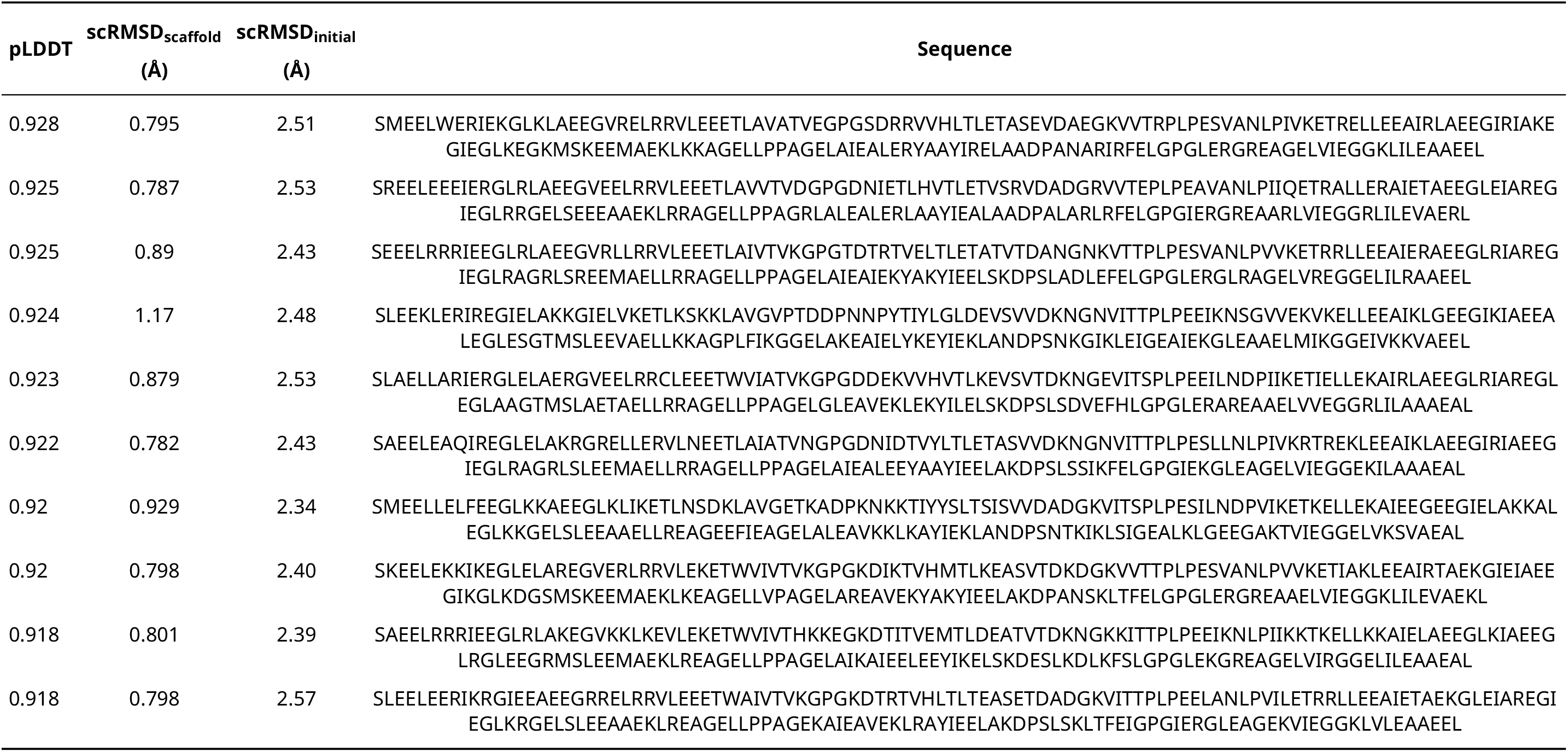
List of Design Sequences for Protein 4p79’s Soluble Analogues.

**Table S6:**
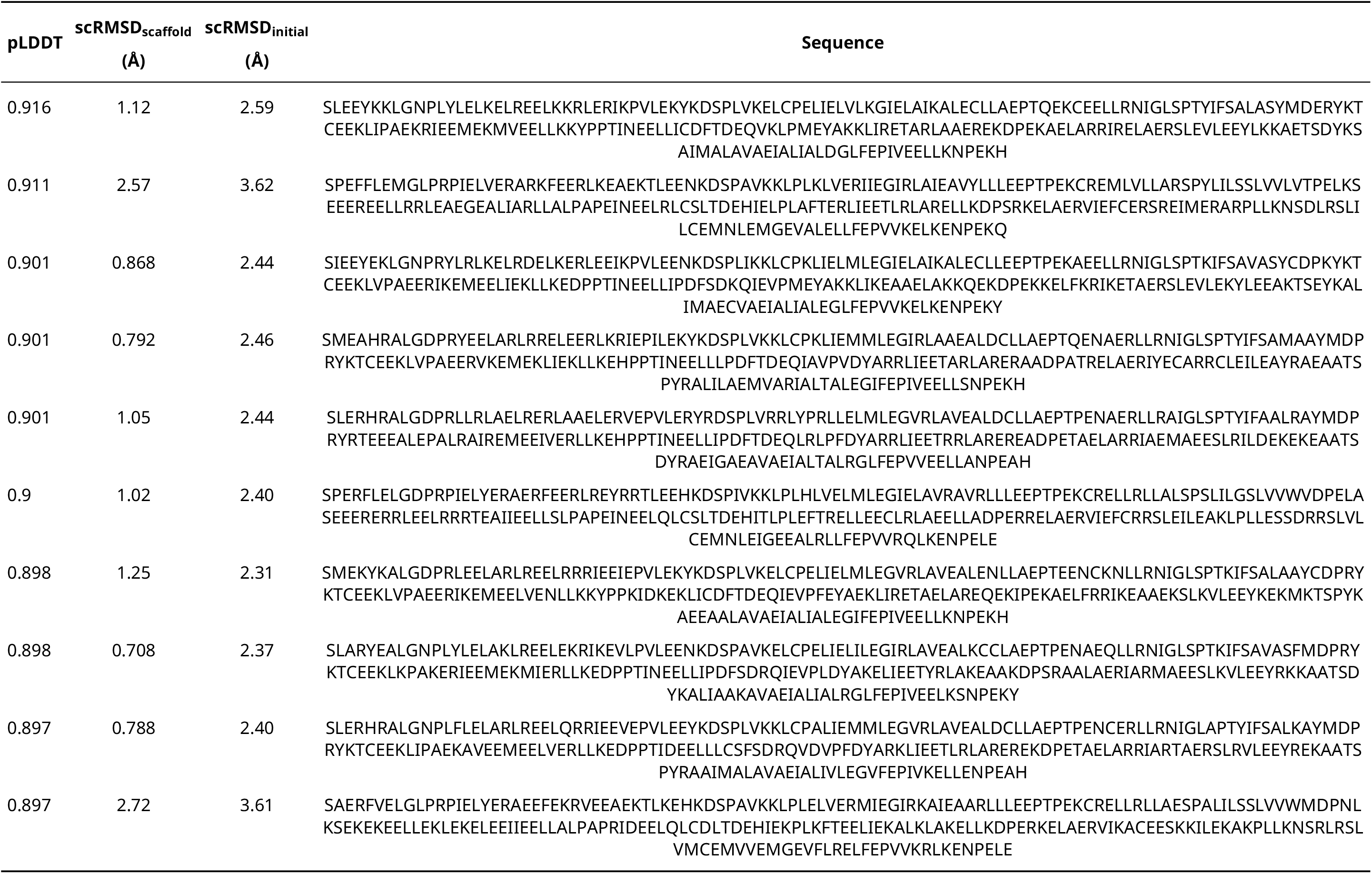
List of Design Sequences for Protein 6ffi’s Soluble Analogues.

### 3 Supplementary Figure

**Figure S1:**
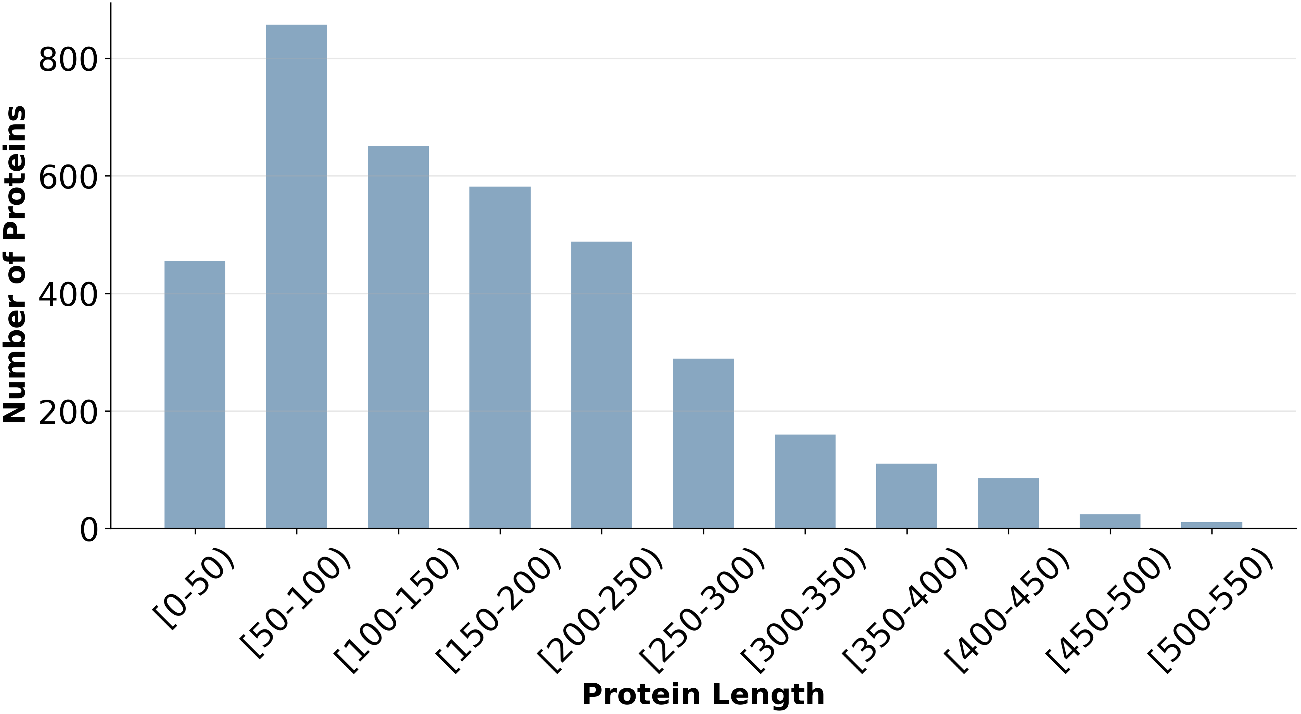
Length distribution of the curated membrane protein domain dataset. The curated dataset for fine-tuning MemProtMPNN comprises 85,051 transmembrane domains extracted from the tmAFDB and filtered to ensure at least 20% transmembrane residues and chain lengths of at least 32 residues, exhibiting a length distribution with a peak around 50–200 residues.

**Figure S2:**
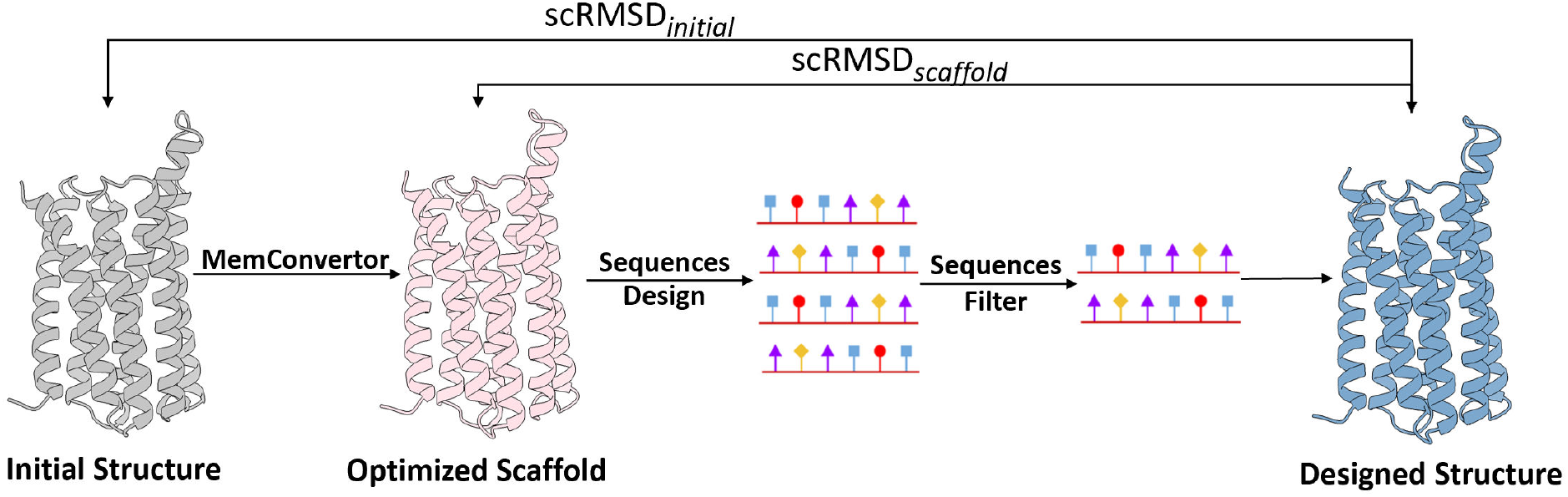
Schematic overview of the MemConverter pipeline and structural consistency metrics. scRMSD_scaffold_ measures C_*α*_ RMSD between the optimized scaffold and designed structure, indicating backbone folding fidelity. scRMSD_initial_ quantifies C_*α*_ RMSD between the initial and designed structures, reflecting core topology preservation.

